# Jasmonate-responsive group IX AP2/ERF transcription factors control the biosynthesis of benzylisoquinoline alkaloids

**DOI:** 10.64898/2026.08.30.748054

**Authors:** Yasuyuki Yamada, Yasuki Tatsumi, Akiko Inagaki, Nobukazu Shitan, Fumihiko Sato

**Author notes:** **Corresponding Authors:** Yasuyuki Yamada, Laboratory of Medicinal Cell Biology, Kobe Pharmaceutical University, 4-19-1 Motoyamakita-machi, Higashinada-ku, Kobe 658-8558, Japan, The author responsible for distribution of materials integra to the findings presented in this article in accordance with the policy described in the Instructions for Authors (https://academic.oup.com/plcell/pages/General-Instructions) is: Yasuyuki Yamada, Fumihiko Sato, Bioorganic Research Institute, Suntory Foundation for Life Science, 8-1-1 Seikadai, Seika-cho, Soraku-gun, Kyoto 619-0284, Japan.

## Abstract

Although the biosynthetic pathways of benzylisoquinoline alkaloids (BIAs) have been extensively investigated in several plant species, their transcriptional regulatory mechanisms remain only partially understood. Jasmonate (JA)-responsive group IX APETALA2/Ethylene Responsive Factor (AP2/ERF) transcription factors (TFs) are well-known regulators of specialized plant metabolism, including the biosynthesis of various alkaloids. However, their specific roles in BIA biosynthesis remain largely elusive. Here, we isolated five novel group IX AP2/ERF TFs, designated Benzylisoquinoline alkaloid Jasmonate-responsive AP2/ERF (BJE1–5), from *Coptis japonica*. Phylogenetic analysis revealed that Benzylisoquinoline alkaloid Jasmonate-responsive AP2/ERF (BJE) proteins belong to subclades distinct from group IXa, which contains well-known AP2/ERF TFs involved in alkaloid biosynthesis. Transient expression analyses in *C. japonica* protoplasts demonstrated that certain BJEs, particularly CjBJE3 and CjBJE5, positively regulated BIA biosynthetic genes through a mutual regulatory network among *BJE* members. Moreover, *CjBJE3* expression was regulated by CjbHLH1, a unique-type basic helix-loop-helix (bHLH) TF specific to BIA-producing plants. Furthermore, heterologous expression of *CjBJE3* and *CjBJE5* in cultured *Eschscholzia californica* cells significantly enhanced the overall BIA production, particularly by increasing end-product benzophenanthridine BIAs, highlighting several uncharacterized biosynthetic genes clustered in the genome. Our findings suggest that BIA-producing species have developed a specific regulatory network comprised of CjbHLH1 and BJE TFs, providing valuable clues for identifying novel biosynthetic enzymes.

## Introduction

Benzylisoquinoline alkaloids (BIAs) are a diverse group of plant-specialized metabolites, including the analgesics morphine and codeine, the antitumor agent noscapine, and the antimicrobial agents sanguinarine and berberine (Facchini 2001). These pharmaceutically important BIAs are primarily found in several plant families such as Papaveraceae, Ranunculaceae, Berberidaceae, and Menispermaceae; however, their endogenous contents in plants are relatively low. Metabolic engineering holds great potential for improved production of valuable BIAs. However, its success is heavily dependent on a deep understanding of the biosynthetic pathways and regulatory mechanisms that orchestrate the expression of biosynthetic enzyme genes.

In recent years, significant progress has been made in identifying and characterizing genes encoding biosynthetic enzymes for berberine in *Coptis japonica* and *Coptis chinensis* (Ranunculaceae), sanguinarine in *Eschscholzia californica* (Papaveraceae), and morphine and noscapine in *Papaver somniferum* (Papaveraceae) (Sato 2013, 2020; Singh et al. 2019; Liu et al. 2021; Becker et al. 2023). Conversely, the transcription factors (TFs) that regulate BIA biosynthesis remain poorly characterized (Yamada and Sato 2013, 2021). CjWRKY1 was isolated from *C. japonica* as the first TF involved in BIA biosynthesis and is reportedly post-transcriptionally regulated (Kato et al. 2007; Yamada and Sato 2016). Subsequently, PsWRKY has been confirmed to be involved in the regulation of morphine biosynthesis (Mishra et al. 2013). Furthermore, a unique type of basic helix-loop-helix (bHLH) TF, CjbHLH1, and its homologs, EcbHLH1-1 and EcbHLH1-2, were isolated and characterized as general transcriptional activators of BIA biosynthesis (Yamada et al. 2011a, 2011b, 2015). Proteins that are highly homologous to CjbHLH1 are exclusively found in BIA-producing species (Yamada et al. 2011a), implying the presence of a specific regulatory system mediated by CjbHLH1 homologs in BIA biosynthesis. However, metabolic engineering using TFs to increase alkaloid productivity remains extremely limited, owing to a lack of detailed information on complicated regulatory systems. Although the ectopic expression of *Arabidopsis thaliana* WRKY1 (AtWRKY1), *E. californica* and *P. somniferum* or CjWRKY1 in cultured *E. californica* cells reportedly increases the accumulation of several BIAs (Apuya et al. 2008; Yamada et al. 2017), studies utilizing TFs for alkaloid engineering remain scarce.

To elucidate the detailed regulatory mechanisms and investigate the transcriptional networks involved in BIA biosynthesis, we previously characterized the *cis*-acting elements in the promoter regions of biosynthetic enzyme genes using cultured *C. japonica* cells (Yamada et al. 2016). Our analysis revealed that a GCC-box-like element, a potential target for the plant-specific APETALA2/Ethylene Responsive Factor (AP2/ERF) family of TFs, is likely involved in transcriptional activation. Furthermore, the ectopic expression of *A. thaliana* and soybean AP2/ERF TFs in *E. californica* and *P. somniferum* increased the expression of several BIA biosynthetic enzyme genes, affecting BIA productivity (Apuya et al. 2008). These findings strongly suggest that endogenous AP2/ERF TFs function as key regulators of BIA biosynthesis.

The AP2/ERF superfamily is one of the largest groups of plant-specific TFs, and all its members possess a conserved AP2/ERF domain of approximately 60 amino acids, which is required for DNA binding. The AP2/ERF superfamily is divided into three subfamilies: AP2 (containing two AP2/ERF domains), RAV (containing single AP2/ERF and B3 domains), and ERF (containing a single AP2/ERF domain) (Nakano et al. 2006). The ERF family is further divided into DREB and ERF subfamilies, which play crucial roles in drought tolerance and various stress responses, respectively (Mizoi et al. 2012; Licausi et al. 2013). Extensive studies have revealed the pivotal roles of AP2/ERF TFs in the regulation of specialized plant metabolism, particularly in members of the jasmonate (JA)-responsive group IX subfamily. Examples include octadecanoid derivative-responsive Catharanthus AP2-domain (ORCA) 2-6 from *Catharanthus roseus* regulating moterpenoid indole alkaloid (MIA) biosynthesis (Menke et al. 1999; van der Fits and Memelink 2000; Paul et al. 2017, 2020; Singh et al. 2020), NtERF189 and NtERF221/ORC1 from *Nicotiana tabacum* regulating nicotine biosynthesis (Shoji et al. 2010; De Boer et al. 2011), GLYCOALKALOID METABOLISM (GAME) 9/JA-responsive ERF (JRE) 4 from *Solanum lycopersicum* regulating steroidal glycoalkaloid (SGA) biosynthesis (Cárdenas et al. 2016; Thagun et al. 2016; Nakayasu et al. 2018), OpERF2 from *Ophiorrhiza pumila* regulating MIA biosynthesis (Udomsom et al. 2016), and AaERF1, AaERF2 and AaORA from *Artemisia annua* regulating artemisinin biosynthesis (Yu et al. 2012; Lu et al. 2013). Group IX AP2/ERF TFs often coordinate with the core JA-signalling complex containing CORONATINE INSENSITIVE 1 (COI1), Jasmonate ZIM domain (JAZ), and MYC2 (De Geyter et al. 2012; Yamada and Sato 2021).

Despite their importance in other specialized metabolic pathways, no group IX AP2/ERF TFs that are definitively involved in the regulation of BIA biosynthesis have been identified to date. Although *C. chinensis* AP2/ERF TFs were recently reported to control gene expression during BIA biosynthesis, they belong to the DREB subfamily (Zhang et al. 2024). In a previous study, we identified 134 AP2/ERF genes in the *E. californica* genome, of which four MeJA-responsive group IX AP2/ERF genes were selected as candidates (Yamada et al. 2020). Although genomic information on *E. californica* is now available, the lack of an efficient transient assay system remains a limiting factor for rapid functional analyses. In contrast, we previously demonstrated that the protoplast-based transient expression system of *C. japonica* is highly advantageous for the functional characterization of TFs (Yamada et al. 2010).

In this study, we identified and characterized group IX AP2/ERF genes from *C. japonica* due to the advantageous analytical platform. We investigated their functions through transient RNAi and overexpression in *C. japonica* protoplasts and assessed their DNA-binding activities using dual-luciferase (LUC) reporter and electromobility shift assays (EMSA). Furthermore, we carried out heterologous expression of AP2/ERF genes in cultured *E. californica* cells to evaluate their effects on biosynthetic gene expression and BIA productivity in stable transformants. Our findings not only contribute to the discovery of novel BIA biosynthetic enzyme genes, but also provide significant insights into the evolution of regulatory networks in specialized plant metabolism.

## Results

### Isolation of MeJA-responsive AP2/ERF TF genes from C. japonica

Based on our previous promoter analyses of berberine biosynthetic genes (Yamada et al. 2016), we explored AP2/ERF candidate genes in the expressed sequence tag (EST) data of *C. japonica*. However, transient RNAi screening of the six candidate sequences did not affect the expression of the *6OMT* gene (Figure S1). Therefore, we isolated new AP2/ERF TF genes using degenerate PCR with cDNAs derived from MeJA-treated cultured *C. japonica* cells. This yielded five sequences, which we designated as “Benzylisoquinoline alkaloid Jasmonate-responsive AP2/ERF 1-5” (CjBJE1-5). Comparison of amino acid sequences of CjBJE1-5 revealed that CjBJE4 and CjBJE5 shared 67.12% identity, whereas all other pairs shared less than 50% identity (Figure S2). Furthermore, CjBJE1-5 was similar to other group IX AP2/ERF TFs involved in the biosynthesis of nicotine (NtERF189), MIA (ORCAs and OpERF2), SGA (GAME9/JRE4), and artemisinin (AaERF1 and AaORA). A phylogenetic tree based on the ERF domain also indicated that CjBJE1 belonged to group IXa, with NtERF189, ORCAs, and GAME9/JRE4; CjBJE2 and CjBJE3 belonged to group IXb; and CjBJE4 and CjBJE5 belonged to group IXc (Figure 1A). Consistent with previous findings that group IX AP2/ERF genes are induced by MeJA, the five *CjBJE* genes exhibited apparent MeJA responsiveness (Figure 1B).

**Figure 1.**
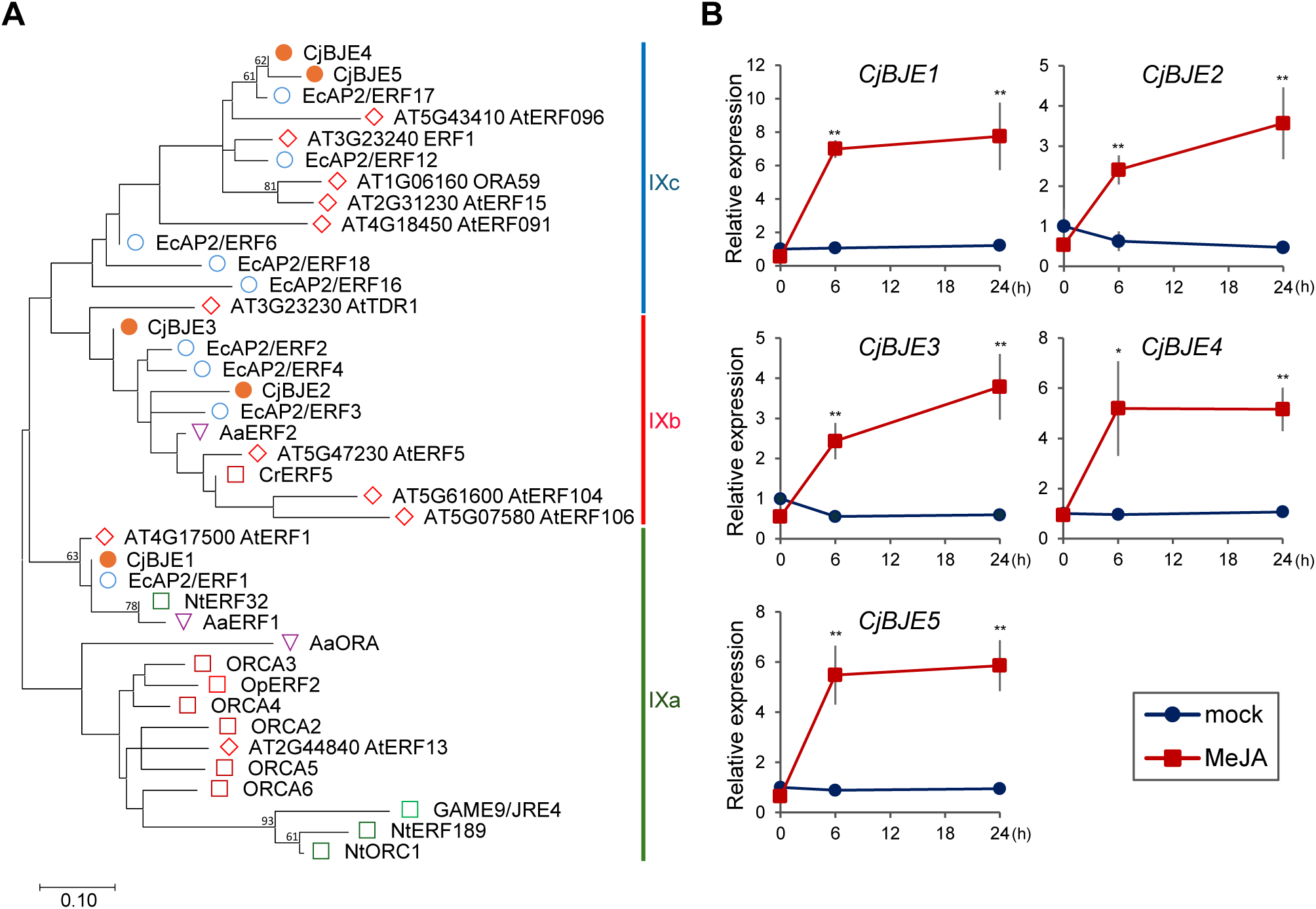
Identification of genes encoding group IX AP2/ERF transcription factors from *Coptis japonica*. (A) Phylogenetic relationships of CjBJE proteins with other characterized group IX AP2/ERFs including sequences from *Arabidopsis thaliana* (At), *Eschscholzia californica* (Ec), *Catharanthus roseus* (Cr), *Nicotiana tabacum* (Nt), *Artemisia annua* (Aa), *Solanum lycopersicum*, and *Ophiorrhiza pumila* (Op). ML tree was constructed from the amino acid sequences of the AP2/ERF domain using MEGA 12. Numbers at nodes indicate bootstrap support values (% from 1,000 replicates); values lower than 50% are omitted for clarity. (B) Expression levels of *CjBJE* genes in cultured *C. japonica* cells treated with methyl jasmonate. Transcript levels were determined by qRT-PCR and normalized to β*-actin*. Relative expression is shown in terms of fold-change compared to mock 0 h sample (set to 1). Error bars indicate standard deviations calculated from three biological replicates. Asterisks denote significant differences compared with the respective mock samples (Student’s *t*-test): \**p* < 0.05; \*\**p* < 0.01.

In *C. japonica*, berberine and other BIAs primarily accumulate in the rhizomes, whereas their biosynthesis occurs in the roots. Tissue expression analysis revealed that biosynthetic enzyme genes (*CjNMCH*, *Cj4’OMT*, and *CjCAS*) showed the highest expression in roots. In contrast, the expression patterns of *CjBJE1-5* were quite different (Figure S3). The expression levels of *CjBJE1*, *CjBJE4*, and *CjBJE5* were high in leaf blades and roots, whereas those of *CjBJE2*, *CjBJE3*, and *CjbHLH1* were high in roots and rhizomes. We also examined the expression levels of *CjBJEs* in 156-S and CjY cultured cells, which exhibited high and low berberine accumulation, respectively (Figure S4A). The expression of *CjBJE1*, *CjBJE2*, *CjBJE3*, and *CjBJE4* was higher in 156-S cells, whereas that of *CjBJE5* was higher in CjY cells. Regarding absolute transcript levels, *CjBJE1* was the most abundant, followed by *CjBJE2* and *CjBJE3* at moderate levels, in 156-S cells (Figure S4B). In contrast, the absolute expression levels of *CjBJE4* and *CjBJE5* were low in both cell types.

Subcellular localization analysis revealed that each CjBJE-sGFP fusion protein was localized to the nucleus in 156-S protoplasts. This suggests that CjBJE proteins function as TFs within the nucleus (Figure S5).

### Characterization of the transcriptional regulation activity of CjBJEs on berberine biosynthetic genes using transient

*RNAi* To assess the regulatory activities of CjBJE1-5 on the expression of berberine biosynthetic enzyme genes, we performed transient RNAi screening for each *CjBJE* gene and monitored changes in *Cj6OMT* expression levels. Although suppression of each *CjBJE* gene was confirmed (Figure S6A), the expression of *Cj6OMT* was not affected in *CjBJE1*-, *CjBJE2*-, or *CjBJE4*-RNAi cells. In contrast, *Cj6OMT* expression in *CjBJE3*-RNAi cells was significantly reduced by 52% compared to the control (Figure S6B). Although *Cj6OMT* expression decreased to 76% of the control level in *CjBJE5*-RNAi cells, no statistically significant reduction was observed. Based on these results, we focused on *CjBJE3* and *CjBJE5* for further analysis, and examined the expression levels of other berberine biosynthetic genes (Figure 2A).

**Figure 2.**
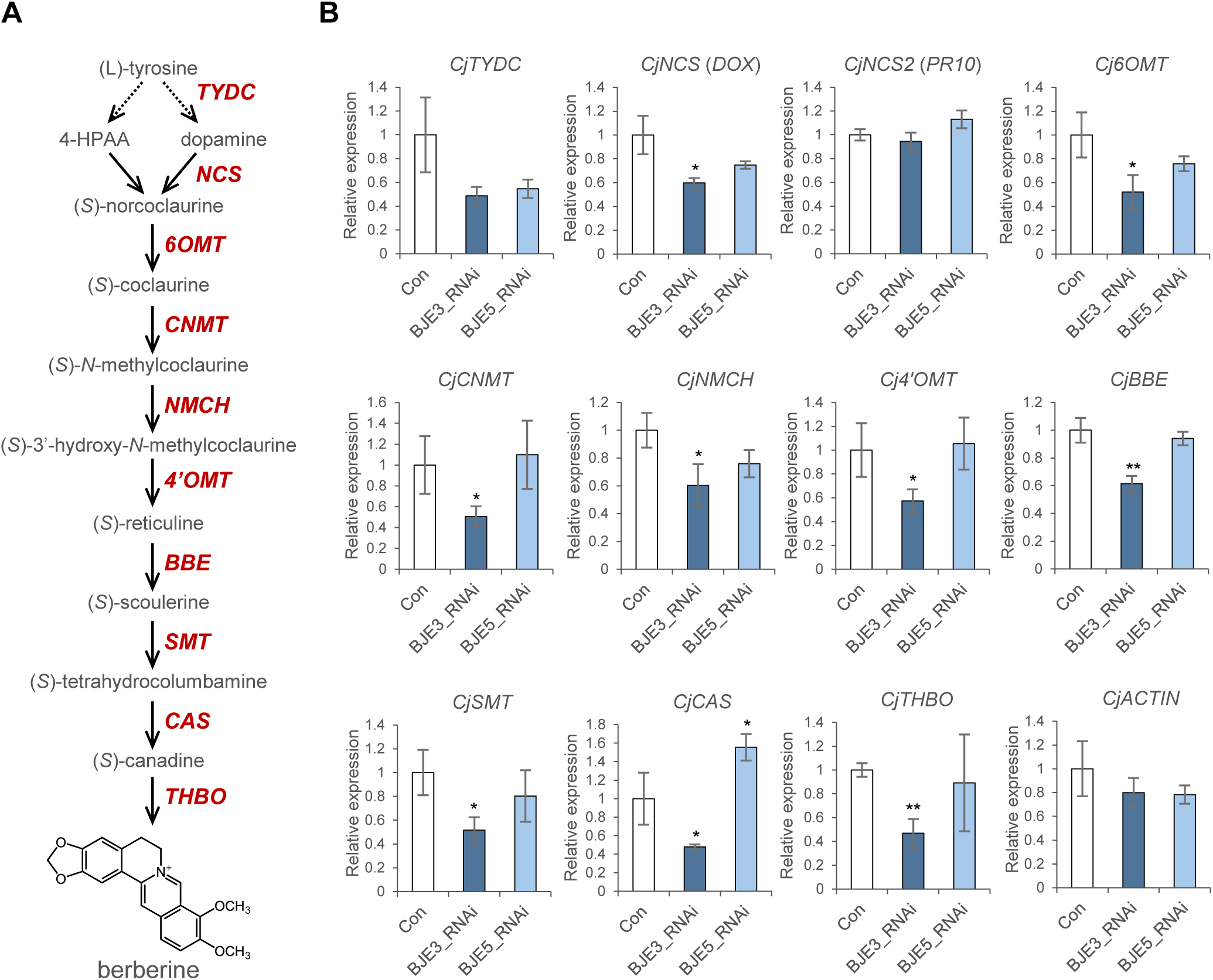
Effect of *CjBJE3* and *CjBJE5* RNA-silencing on the expression of biosynthetic enzyme genes in berberine biosynthesis. (A) Schematic representation of the berberine biosynthetic pathway. Enzymes characterized to date are highlighted in red. Dotted arrows indicate unidentified or uncharacterized steps in the pathway. (B) Transcript levels of berberine biosynthetic enzyme genes and β*-actin* in *C. japonica* protoplasts. Expression levels were determined by qRT-PCR and normalized to the α*-tubulin* gene. Relative expression levels are shown in terms of fold-change compared to control (set to 1). Error bars indicate standard deviations from three biological replicates. Asterisks denote significant differences from the control according to Student’s *t*-test: \**p* < 0.05; \*\**p* < 0.01.

In *CjBJE3*-RNAi cells, the expression levels of almost all berberine biosynthetic enzyme genes were significantly reduced (to 47–61% of the control); however, the decrease in *CjTYDC* and *CjNCS2* (pathogenesis-related protein 10; PR10-type) did not reach the level of statistical significance. In contrast, the expression of almost all the biosynthetic genes remained quite unaffected in *CjBJE5*-RNAi cells. However, the transcript levels of *CjTYDC*, *CjNCS* (dioxygenase; DOX-type), *CjNMCH*, and *CjSMT* in *CjBJE5*-RNAi cells were relatively low, ranging from 55% to 80% of the control levels (Figure 2A). *CjACTIN* expression remained constant in both *CjBJE3*- and *CjBJE5*-RNAi cells, suggesting specific effects of *CjBJE3*- and *CjBJE5*-RNAi on the expression of berberine biosynthetic enzyme genes. These results indicated that CjBJE3 acts as a primary regulator of the berberine biosynthetic pathway, whereas CjBJE5 may play a partial role in its regulation. We also examined the effects of *CjBJE3*- and *CjBJE5*-RNAi on the expression levels of other TF genes. In both RNAi lines, the expression levels of *CjbHLH1* and *CjWRKY1* were not significantly altered, although they decreased to 82% and 62% of the control levels, respectively, in *CjBJE3*-RNAi cells (Figure 3A). In contrast, the transcript levels of other *CjBJE* genes were significantly reduced to 34–58% of the control levels in both *CjBJE3*- and *CjBJE5*-RNAi cells (Figure 3B). These results suggest that CjBJE3 and CjBJE5 regulate the expression of other *CjBJE* genes, indicating the presence of a regulatory network among CjBJEs.

**Figure 3.**
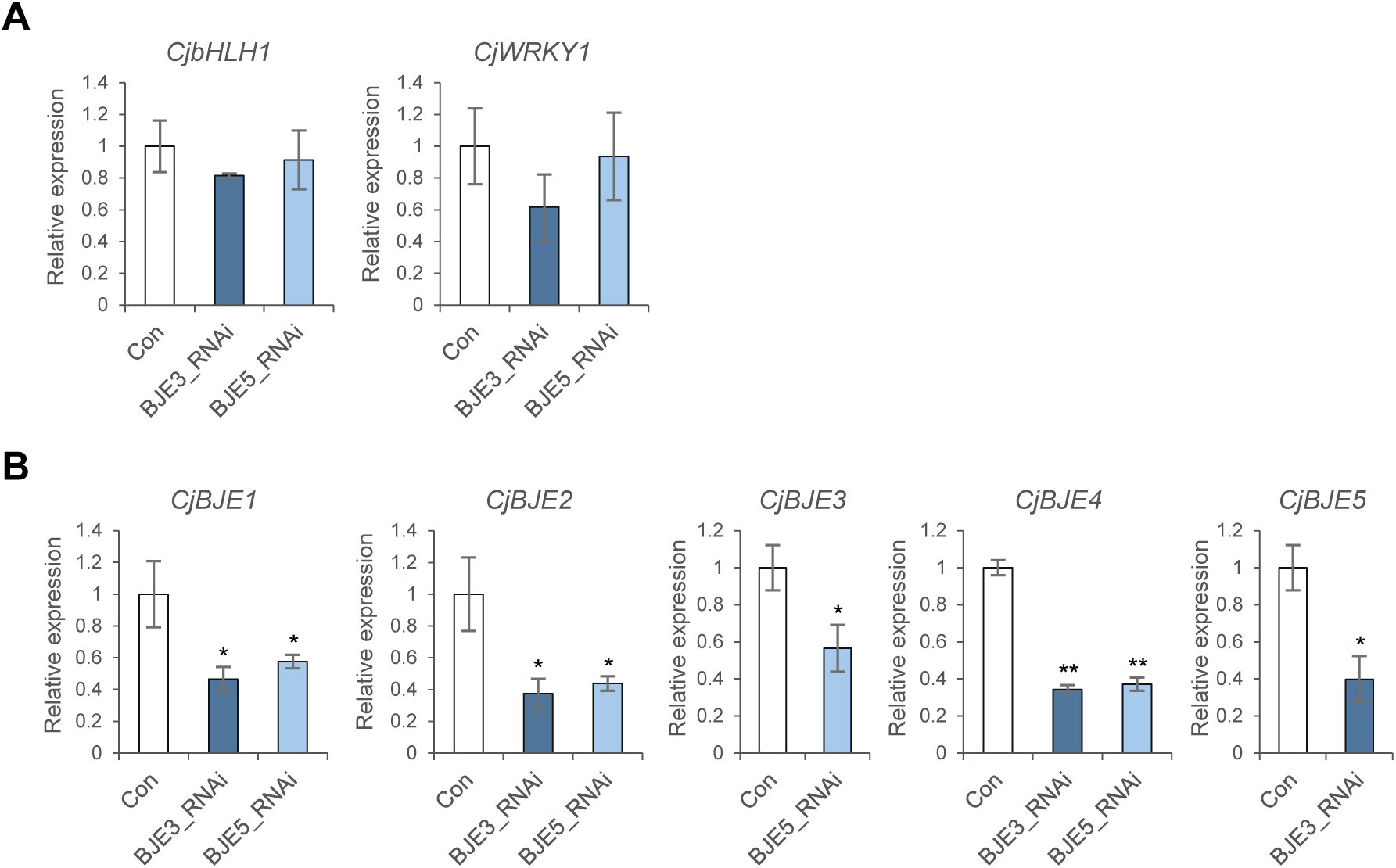
Effect of *CjBJE3* and *CjBJE5* RNA-silencing on the expression of transcription factor genes. Transcript levels of *CjbHLH1*, *CjWRKY1* (A), and other *CjBJE* genes (B) in *C. japonica* protoplasts. Expression levels were determined by qRT-PCR and normalized to α*-tubulin*. Relative expression levels are shown in terms of fold-change compared to control (set to 1). Error bars indicate the standard deviation of three biological replicates. Asterisks denote significant differences from control according to Student’s *t*-test: \**p* < 0.05; \*\**p* < 0.01.

### Transactivation activities of CjBJE3 and CjBJE5 on berberine biosynthesis genes using transient overexpression

Transient RNA interference (RNAi) analysis suggested that CjBJE3 and CjBJE5 act as transcriptional activators of berberine biosynthesis. To further validate this hypothesis, we examined the effects of overexpressing *CjBJE3* and *CjBJE5* in 156-S protoplasts. The transcript levels of most berberine biosynthetic enzyme genes significantly increased in both *CjBJE3*-OX and *CjBJE5*-OX cells (Figure 4A). In *CjBJE3*-OX cells, the expression levels of *CjTYDC*, *CjNCS2* (PR10-type), *Cj6OMT*, *CjNMCH*, *Cj4’OMT*, *CjBBE*, and *CjCAS* were significantly upregulated, ranging from 1.2- to 5.1-fold that of the control. Similarly, in *CjBJE5*-OX cells, the expression of *CjTYDC*, *CjNCS2* (PR10-type), *Cj6OMT*, *CjNMCH*, *Cj4’OMT*, and *CjCAS* was significantly induced, by 1.5- to 4.9-fold. In contrast, the expression of *CjNCS* (DOX-type), *CjCNMT*, *CjSMT*, and *CjTHBO* was either reduced or remained unchanged in both cell lines. When we examined the effects of *CjBJE3*- and *CjBJE5*-overexpression on other TF genes, the expression of *CjbHLH1* and *CjWRKY1* was significantly upregulated by 2.5-fold in *CjBJE3*-OX and 3.0-fold in *CjBJE5*-OX cells, respectively (Figure 4B). In particular, the overexpression of *CjBJE3* led to a significant increase in the expression of *CjBJE2*, *CjBJE3*, and *CjBJE4*, whereas the overexpression of *CjBJE5* significantly decreased their expression (Figure 4B). These results confirm that CjBJE3 and CjBJE5 function as transcriptional activators of berberine biosynthesis and suggest the presence of a complex regulatory network involving *CjbHLH1*, *CjWRKY1*, and other AP2/ERF TFs.

**Figure 4.**
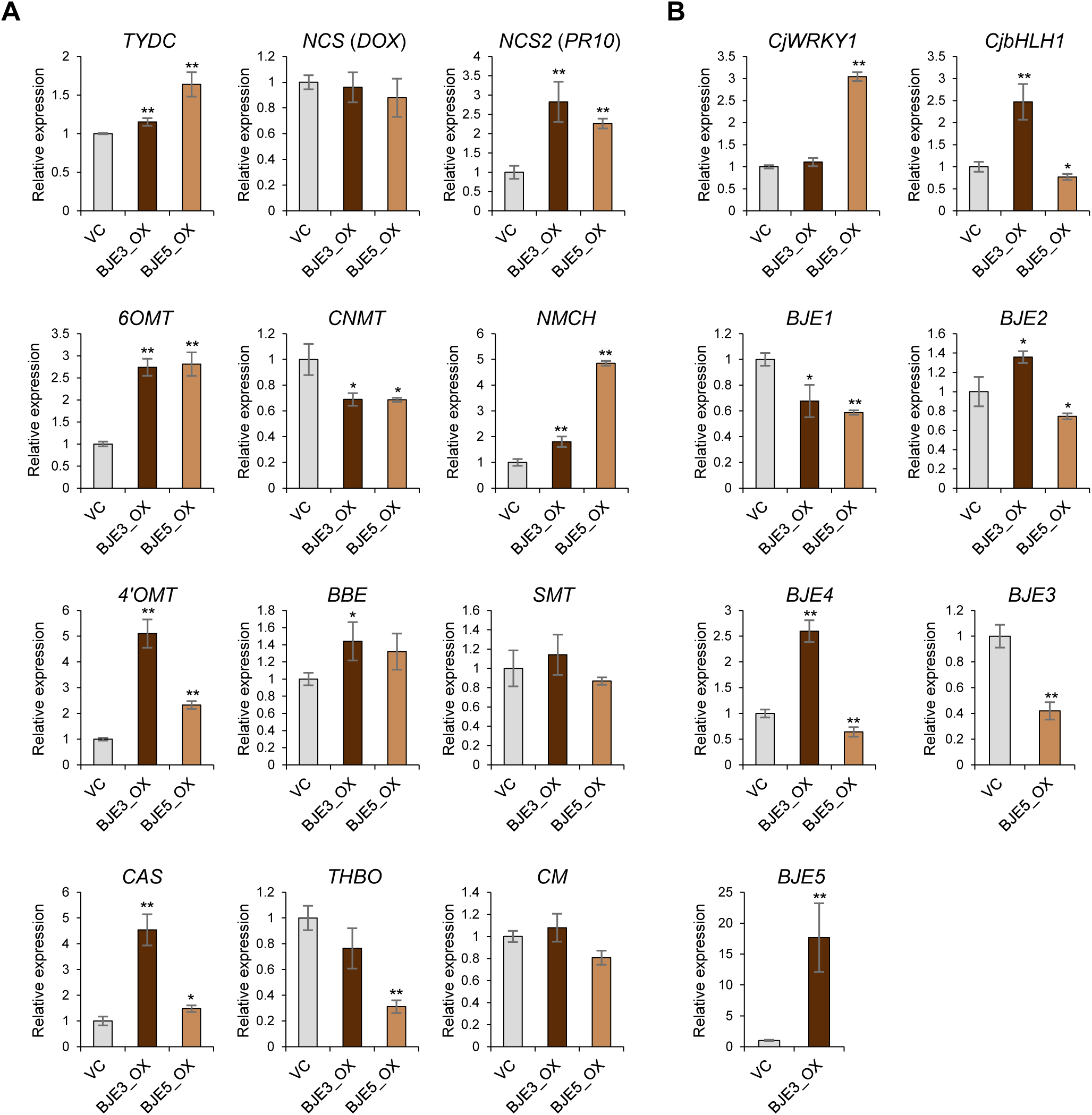
Effect of *CjBJE3* and *CjBJE5* overexpression on the expression of genes involved in berberine biosynthesis. Transcript levels of berberine biosynthetic enzyme (A) and transcription factor (B) genes in *C. japonica* protoplasts. Expression levels were determined by qRT-PCR and normalized to α*-tubulin*. Relative expression levels are shown in terms of fold-change compared to control (set to 1). Error bars indicate standard deviation of three biological replicates. Asterisks denote significant differences from control according to Student’s *t*-test: \**p* < 0.05; \*\**p* < 0.01.

Our previous promoter analyses of *Cj4’OMT* and *CjCAS* revealed that a putative GCC-like *cis*-element is essential for their activation. To confirm the direct interaction between these promoters and CjBJE3 or CjBJE5, we performed an EMSA using the purified GST-fused CjBJE3 and CjBJE5 proteins. Since the full-length CjBJE3 protein failed to stably express in *Escherichia coli* cells (Figure S7), an N-terminal deletion of 61 amino acids (hereafter referred to as GST-CjBJE3Δ61) was introduced. Shifted bands were detected when the DNA probe was incubated with GST-CjBJE3Δ61 or GST-CjBJE5 (Figure S8). Notably, the shifted bands for GST-CjBJE5 were more distinct than those for GST-CjBJE3Δ61, suggesting a higher binding affinity of GST-CjBJE5. The appearance of broad or multiple-shifted bands was likely due to the instability of the purified proteins, which showed minor degradation products even on Coomassie brilliant blue-stained polyacrylamide gels (Figure S7). These DNA–protein complexes disappeared upon the addition of an excess of unlabeled competitors containing the GCC-like sequence.

To further investigate the transactivation capacity of CjBJE3 and CjBJE5 via this GCC-like sequence, dual-LUC reporter assays were performed in 156-S protoplasts using reporter vectors driven by either native (AGCCACC) or mutated (AGTAACT; mGCC) promoters (Figure 5A). The relative LUC activities driven by the *Cj4’OMT* and *CjCAS* promoters were markedly induced by both CjBJE3 and CjBJE5; however, this induction was significantly attenuated when the mGCC promoters were used (Figure 5B). Notably, CjBJE5 exhibited a more drastic decrease in LUC reporter activity on the mutated promoters than CjBJE3, which was consistent with the EMSA results. These results indicate that CjBJE3 and CjBJE5 directly bind to the promoter regions of *Cj4’OMT* and *CjCAS* to regulate their expression. Although the mutation of the GCC-like sequence significantly attenuated promoter activity, both CjBJE3 and CjBJE5 displayed residual transactivation capabilities, suggesting the existence of additional non-canonical *cis*-acting elements within these promoter regions.

**Figure 5.**
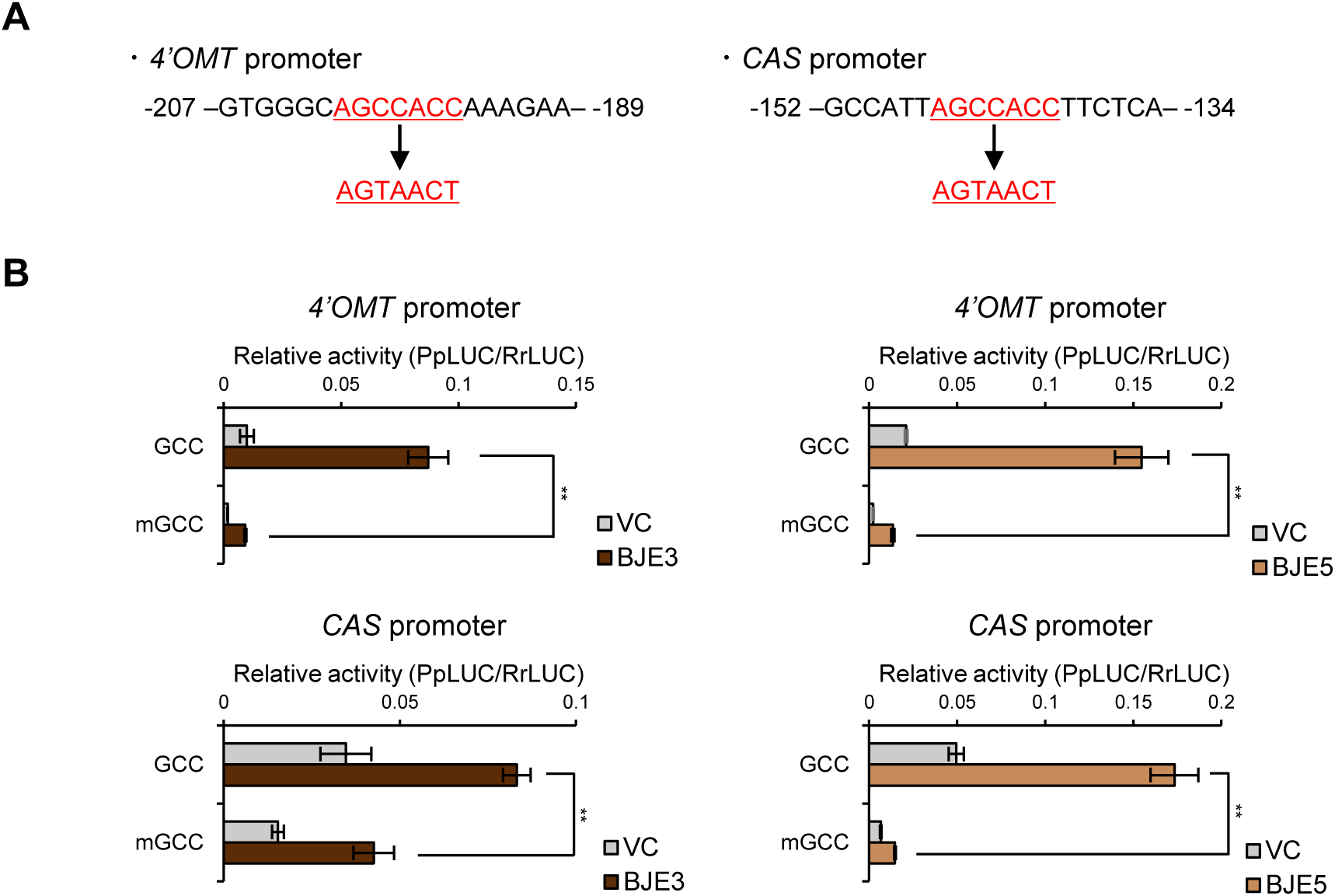
Binding activity of CjBJE3 and CjBJE5 to the promoter region of *4’OMT* and *CAS* genes. (A) Mutation in GCC-like sequence (AGCCACC) in promoter region of *4’OMT* and *CAS* genes. (B) Transactivation of promoter activities by CjBJE3 and CjBJE5 in *C. japonica* protoplasts. Promoter activities were evaluated using a transient dual-LUC reporter assay in *C. japonica* protoplasts. Firefly luciferase activity was normalized to *Renilla* luciferase activity as an internal control. Data represent mean ± SD of three independent biological replicates. Asterisks indicate significant differences compared with control (Student’s *t*-test: \**p* < 0.05, \*\**p* < 0.01).

### RNA-silencing and overexpression of CjbHLH1 affects the expression of CjBJE3

Our previous studies revealed that CjbHLH1 and its homologs, a unique type of bHLH TFs found in BIA-producing species, act as important transcriptional activators in *C. japonica* and *E. californica* (Yamada et al. 2011a, 2015). To investigate whether *CjBJE* genes were regulated by CjbHLH1, we performed transient RNAi and overexpressed *CjbHLH1* in 156-S protoplasts. The suppression of *CjbHLH1* significantly decreased the expression of *CjBJE3* to 38% of the control level, a reduction pattern similar to its effect on *CjBJE1* and *Cj6OMT*, whereas the expression levels of *CjBJE2*, *CjBJE4* and *CjBJE5* were not significantly altered, although they decreased to 61%, 57%, and 85% of the control levels, respectively (Figure 6A). In contrast, the overexpression of *CjbHLH1* significantly increased the transcript level of *CjBJE3* by 1.3-fold, similar to the induction of *Cj4’OMT* (1.6-fold), while the expression levels of *CjBJE1*, *CjBJE2*, *CjBJE4*, and *CjBJE5* were not altered or suppressed (Figure 6B). Although the expression of *CjBJE3* increased in *CjbHLH1*-OX cells, the expression of *CjBJE2* and *CjBJE5* showed no significant response, which could be due to the slight degree of *CjBJE3* upregulation. These results indicated that CjbHLH1 regulates berberine biosynthesis by directly activating biosynthetic enzyme genes and indirectly inducing their expression by upregulating *CjBJE3*.

**Figure 6.**
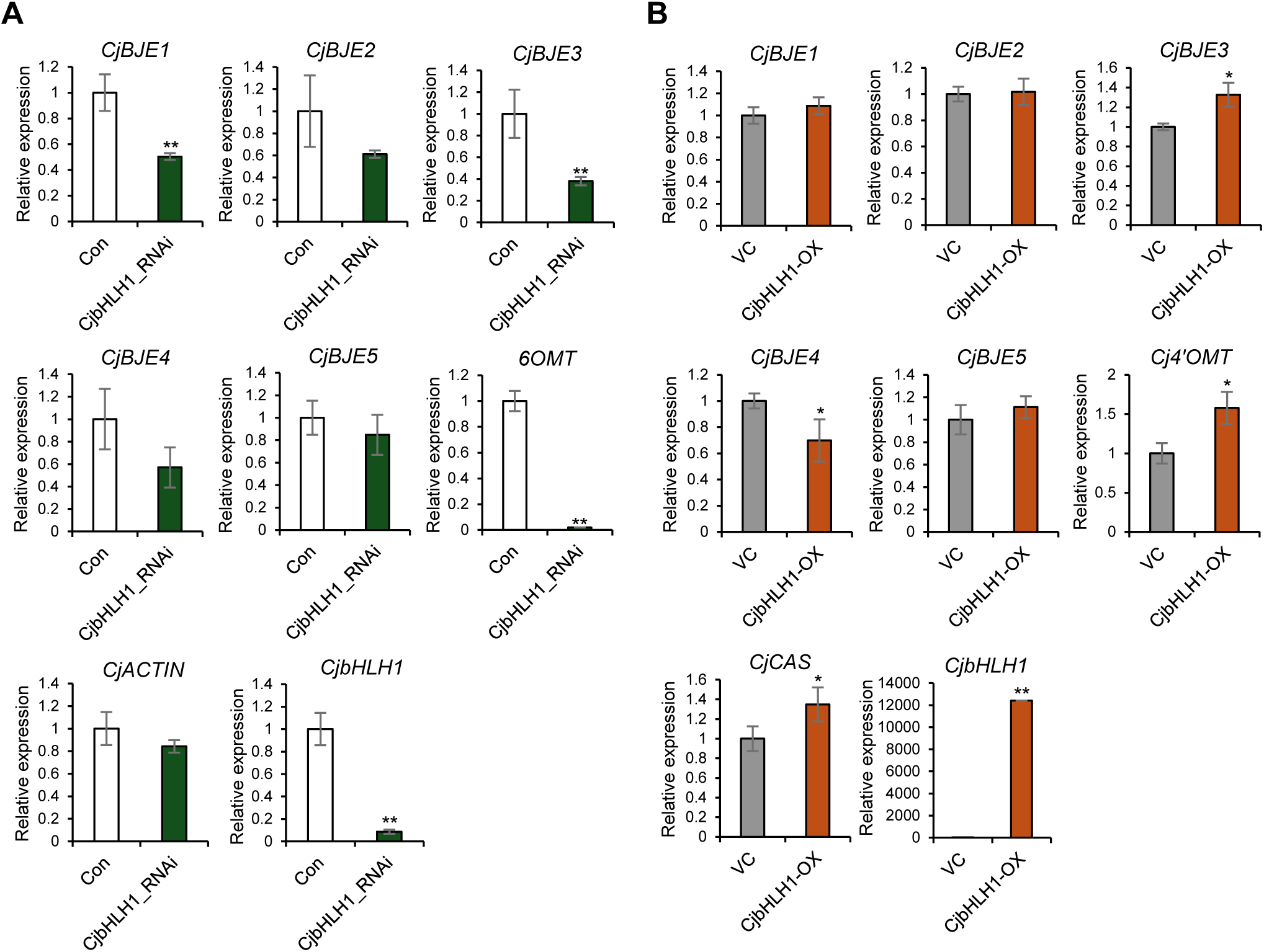
CjbHLH1 regulates the expression of *CjBJEs* in *Coptis japonica*. Effects of *CjbHLH1* RNA-silencing (A) and *CjbHLH1* overexpression (B) on expression of other *CjBJE*, and *Cj6OMT* or *Cj4’OMT* and *CjCAS* genes. Expression levels were determined by qRT-PCR and normalized to the α*-tubulin*. Relative expression levels are shown in terms of fold-change compared to control (set to 1). Error bars indicate standard deviation of three biological replicates. Asterisks denote significant differences from control according to Student’s *t*-test: \**p* < 0.05; \*\**p* < 0.01.

### Heterologous expression of CjBJE3 and CjBJE5 genes in cultured E. californica cells

To assess the functions of CjBJE3 and CjBJE5 in stable transformants, we used *E. californica* (California poppy). *E. californica* shares the biosynthetic pathway from (*S*)-norcoclaurine to (*S*)-scoulerine with *C. japonica* but primarily accumulates benzophenanthridine-type BIAs, such as sanguinarine, chelerythrine, chelirubine, and macarpine, in the roots and cultured cells (Figure 7A). Given the availability of a draft genome sequence for the *E. californica* cultivar ‘Hitoezaki’ and the established stable transformation methods, we performed a heterologous expression analysis in this species (Hori et al. 2018; Yamada et al. 2021). Transgenic *E. californica* cells were generated via *Agrobacterium*-mediated transformation using *CjBJE3*- and *CjBJE5*-overexpression (OX) vectors, with a β-glucuronidase (*GUS*) expression vector as a vector control (VC). In addition, we used the CRES-T method by fusing the SRDX repressor domain with CjBJE3 and CjBJE5 to express chimeric repressors in California poppy cells (Hiratsu et al. 2003). Following the induction of transgenic calli and kanamycin selection, 3–5 transgenic lines were maintained in liquid culture for several months (Figure S9A). After confirming the integration of exogenous genes by genomic PCR (Figure S9B), three VC, three OX, and two SRDX cell lines were selected for gene expression and metabolite analyses. RT-PCR analysis indicated the successful expression of the heterologous genes (Figure S9C). Notably, the growth of *CjBJE3*-OX and *CjBJE5*-OX cell lines was slower than that of the other lines, and these OX lines showed distinct pigmentation in the culture medium (Figure S9A).

**Figure 7.**
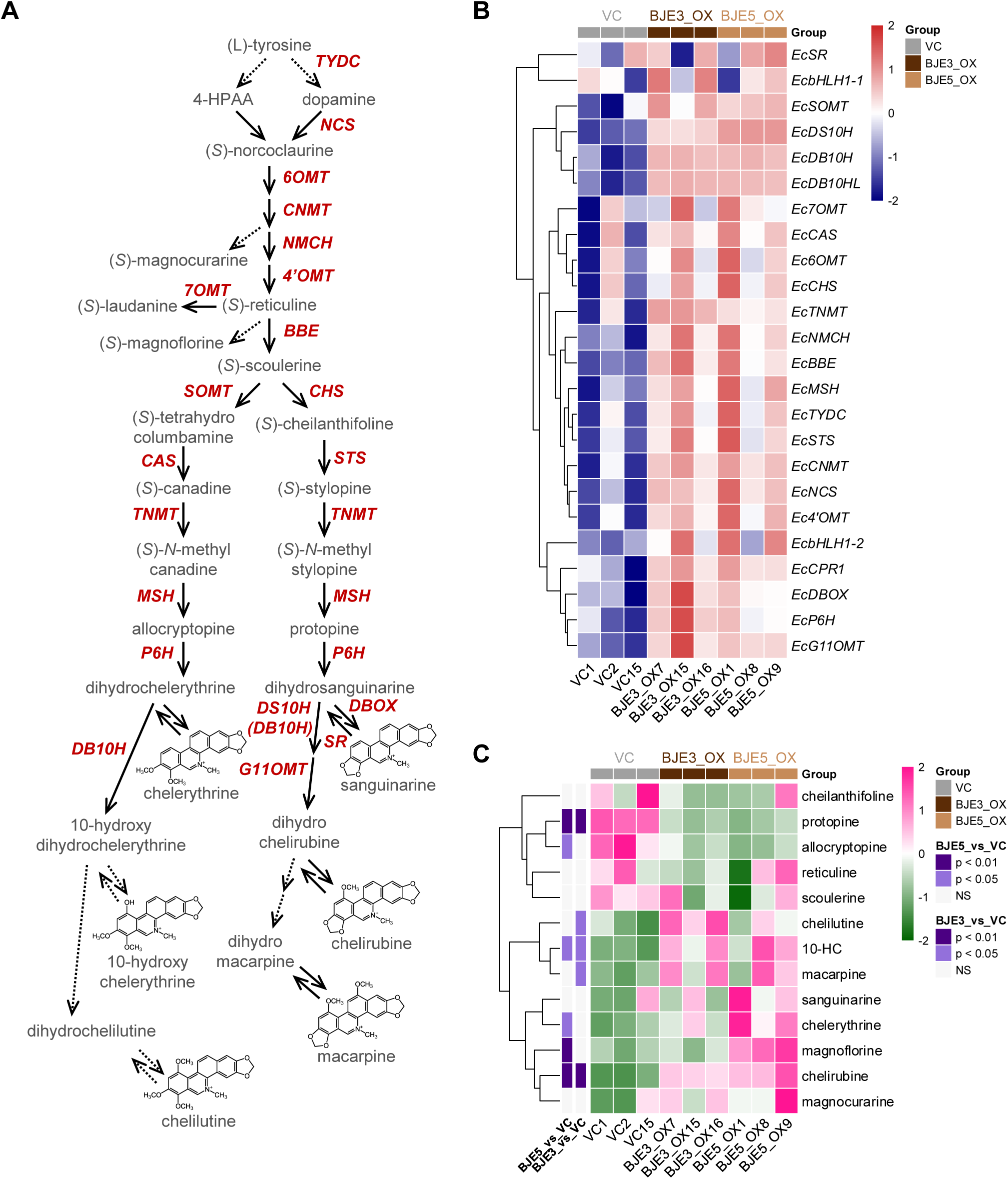
Heterologous expression of *CjBJE3* and *CjBJE5* in cultured *Eschscholzia californica* cells. (A) Biosynthetic pathways of benzylisoquinoline alkaloids found in cultured *E. californica* cells. Enzymes are indicated in red. Dotted arrows represent unidentified or uncharacterized enzymatic steps. (B) Gene expression profiling in transgenic cultured *E. californica* cells. Heatmap and hierarchical clustering illustrate expression patterns of known BIA biosynthetic genes. Transcript levels (TMM values) were normalized using Z-score transformation to visualize relative expression changes across samples. (C) Metabolite profiling in transgenic cultured *E. californica* cells. Heatmap and hierarchical clustering show relative abundance of alkaloids. Data were standardized by calculating Z-scores for each metabolite across all samples. Purple-to-gray scale represents number of standard deviations from mean concentration.

Although the expression of several biosynthetic enzyme genes (*Ec7OMT*, *EcCAS*, *Ec6OMT*, and *EcCHS*) was unexpectedly higher in the VC2 line, the transcript levels of BIA biosynthetic genes were upregulated in both *CjBJE3*-OX and *CjBJE5*-OX cells compared to those in VC cells (Figure 7B). Although the expression of *EcbHLH1-1* and *EcbHLH1-2*, which are homologs of *CjbHLH1*, increased in several OX lines, the values were inconsistent across lines. Notably, the expression of *EcSOMT*, *EcDS10H*, and *EcDB10H* was markedly upregulated in OX cell lines. Additionally, heatmap analysis indicated that *EcCPR1*, which encodes a cytochrome P450 reductase, was upregulated in both OX lines.

We further analyzed the total content of BIAs, mainly found in cultured *E. californica* cells, cultured cells, and media. As shown in Figure 7C, the BIA profiles of *CjBJE3*-OX and *CjBJE5*-OX cells were considerably different from those of VC cells. The accumulation of 10-hydroxychelerythrine (10-HC), chelilutine, chelirubine, and macarpine in *CjBJE3*-OX lines was significantly higher than in the VC lines, showing a marked increase of approximately 5–10-fold. These BIAs represent the end products of the biosynthetic pathway, indicating that CjBJE3 enhances the metabolic flow by regulating BIA biosynthetic genes. In contrast, the levels of intermediates, such as cheilanthifoline, protopine, allocryptopine, reticuline, and scoulerine, were lower in *CjBJE3*-OX cells, although a significant reduction was observed only for protopine. A similar BIA profile was observed in the *CjBJE5*-OX cells. The accumulation of 10-HC, chelerythrine, chelirubine, and magnoflorine in *CjBJE5*-OX lines was significantly higher than in the VC lines (approximately 3- to 10-fold). Sanguinarine and macarpine accumulated at high levels in *CjBJE5*-OX cells, whereas protopine and allocryptopine levels significantly decreased. The effects of *CjBJE3*-SRDX and *CjBJE5*-SRDX expression on the transcript levels of BIA biosynthetic enzyme genes varied among the cell lines; however, a decrease in the expression of several biosynthetic genes was observed in the SRDX lines (Figure S10A). Furthermore, the accumulation of BIA intermediates (such as reticuline, scoulerine, cheilanthifoline, and protopine) was observed in both SRDX lines, whereas the levels of benzophenanthridine-type BIAs such as sanguinarine, chelirubine, macarpine, chelerythrine, 10-HC, and chelilutine remained unchanged or were reduced (Figure S10B). These results suggest that both CjBJE3 and CjBJE5 act as transcriptional activators in *E. californica* stable transgenic cells. Their activity modulates the metabolic flow of BIA biosynthesis; the overexpression of CjBJE3 and CjBJE5 increased the end products while decreasing the intermediates in BIA biosynthesis, whereas the overexpression of their chimeric repressors led to the opposite metabolic profile.

### Overexpression of CjBJE3 and CjBJE5 upregulated the expression of candidate biosynthetic enzyme genes

Since the heterologous expression of *CjBJE3* and *CjBJE5* enhanced BIA production in transgenic cultured *E. californica* cells, we further investigated the expression of genes encoding biosynthetic enzymes as candidates for catalyzing uncharacterized BIA biosynthetic pathways. Our draft genome analysis and recent high-quality genome analysis of *E. californica* revealed an expansion of the *CYP82* gene family (Hori et al. 2018; Rössner et al. 2026), with several genes clustered within the genome; therefore, we focused on these *CYP82* genes. In addition, genes encoding predicted oxidases, reductases, *O*-methyltransferases (OMTs), *N*-methyltransferases (NMTs), and other cytochrome P450 proteins (CYP71, CYP76, CYP84, and so on) exhibited high trimmed mean of M values (TMM). Heatmap analysis, including characterization of enzyme genes, indicated that several *CYP82N* and *CYP82P* subfamily genes, as well as *OMT* and *NMT* genes, were upregulated in both *CjBJE3*-OX and *CjBJE5*-OX cells (Figure 8). These genes may be involved in the biosynthesis of BIAs that were highly accumulated in *CjBJE*-OX cells.

**Figure 8.**
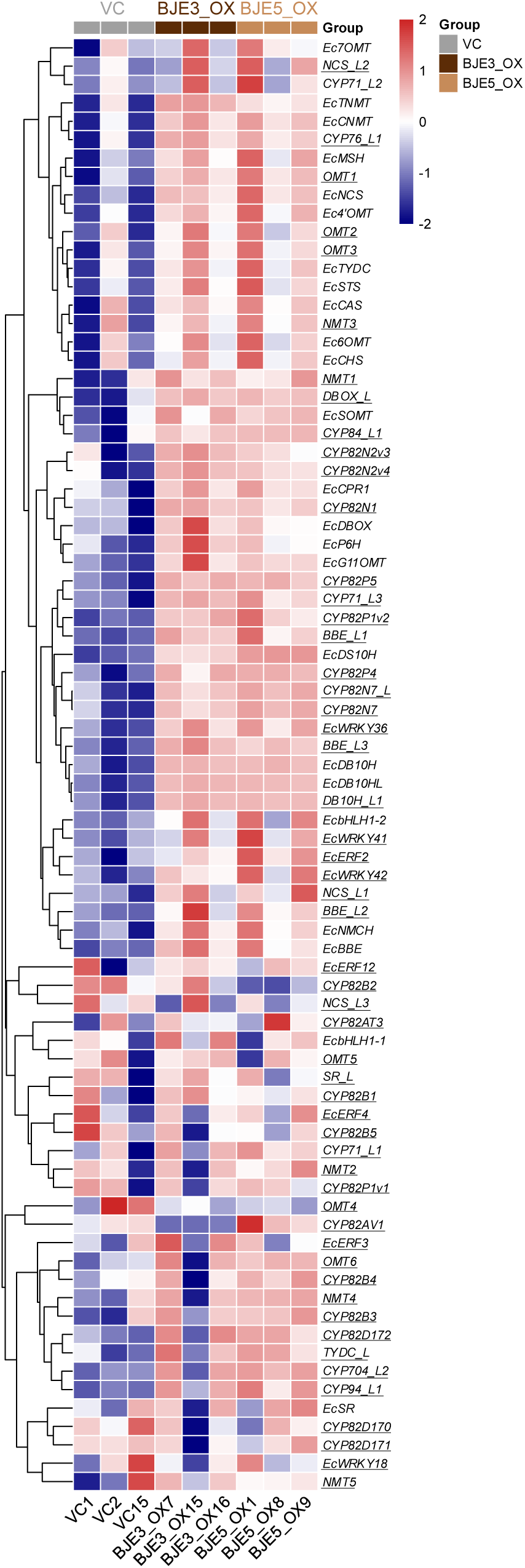
Expression profiling of putative biosynthetic enzyme genes in *CjBJE3*- and *CjBJE5*-overexpressing cultured *E. californica* cells. Heatmap shows relative expression patterns (Z-score transformed TMM values) of genes (underlined) encoding uncharacterized enzymes, including cytochrome P450s (CYPs), *O*-methyltransferases (OMTs), and *N*-methyltransferases (NMTs), along with several transcription factors.

To examine whether these co-regulated genes shared conserved *cis*-elements targeted by CjBJE3 and CjBJE5, we performed MEME motif analysis on the promoter regions of genes markedly upregulated in both *CjBJE3*-OX and *CjBJE5*-OX cells. Although the predicted GCC-like sequence (AGCCACC) found in the *Cj4’OMT* and *CjCAS* promoters was relatively rare, several GC-rich motifs were enriched (Figure S11). Notably, these motifs are conserved in the promoter regions of the clustered *CYP82P* genes, including *DB10H*, *DB10H_L*, *DB10H_L2*, and *CYP82P5*. These results suggest that CjBJE3 and CjBJE5 recognize several variants of the GCC-like sequence and mediate the coordinated transcriptional regulation of pathway genes located in genomic clusters.

## Discussion

As berberine and related BIAs, which accumulate in the rhizomes of *C. japonica*, are pharmaceutically important compounds (Inui et al. 2012), the biosynthetic pathways and regulatory mechanisms of BIAs in *Coptis* species have been extensively investigated using cultured cells and, more recently, genome sequencing (Sato 2013; He et al. 2018; Chen et al. 2021; Yamada and Sato 2021; Liu et al. 2023; Zhang et al. 2024). While a unique bHLH protein, CjbHLH1, is considered to play a crucial role in the transactivation of biosynthetic genes, previous studies have indicated that there is a broader transcriptional network comprising bHLH, WRKY, and AP2/ERF TFs that may be involved in the regulation of BIA biosynthesis (Yamada et al. 2015). In the present study, we identified group IX AP2/ERF TFs, designated BIA jasmonate-responsive AP2/ERFs (BJEs), from *C. japonica* and demonstrated that CjBJE3 and CjBJE5 regulated the expression of BIA biosynthetic enzyme genes in *C. japonica* cells. Heterologous expression of *CjBJE3* and *CjBJE5* in cultured *E. californica* cells upregulated the expression of BIA biosynthetic enzyme genes and enhanced the production of end-products, thereby providing valuable insights for identifying novel candidate genes and putative *cis*-elements involved in BIA biosynthesis.

### Structural diversification and genomic organization of group IX AP2/ERF transcription factors

JA-responsive group IX AP2/ERF TFs regulate the biosynthesis of various specialized metabolites (Yamada and Sato 2021). For instance, in alkaloid biosynthesis, NtERF189 (nicotine biosynthesis in tobacco), ORCAs, OpERF2 (MIA biosynthesis in *C. roseus* and *O. pumila*), and GAME9/JRE4 (SGA biosynthesis in tomato) all belong to the group IXa subclade. In contrast, the results of the present study showed that CjBJE3 and CjBJE5 belong to group IXb and IXc subclades, respectively (Figure 1A). Although CjBJE1 belongs to group IXa, it does not regulate BIA biosynthesis. To date, with regard to alkaloid biosynthesis, only CrERF5 from *C. roseus* has been identified as a positive regulator of group IXb (Pan et al. 2019), and no AP2/ERF proteins from group IXc have been reported as alkaloid regulators. These findings suggest that while group IX AP2/ERF TFs play a conserved role in regulating alkaloid biosynthesis across diverse plant species, including basal eudicots, their protein structures and functions diversified during the evolution of specialized metabolism. This diversification likely led to the formation of specific lineage-dependent regulatory networks.

In many cases, group IX AP2/ERF genes form gene clusters that coordinate the regulation of specialized metabolism (Shoji and Yuan 2021). For instance, specific regulatory genes within the clusters, such as *NtERF189* in tobacco and *GAME9/JRE4* in tomato, encode master regulators of nicotine and SGA biosynthesis, respectively, whereas *ORCA* cluster genes (*ORCA2*–*6*) in *C. roseus* encode AP2/ERF TF proteins that are collectively involved in regulating MIA biosynthesis (Singh et al. 2020). Although it remains to be confirmed whether *CjBJE1–5* forms a gene cluster in the *C. japonica* genome, genomic data from related species provide intriguing insights. In *C. chinensis*, homologs of *CjBJE2* and *CjBJE3* are located on chromosome 6, whereas those of *CjBJE1*, *CjBJE4*, and *CjBJE5* are distributed across different chromosomes (Liu et al. 2021). In addition, in *Aquilegia coerulea*, another member of the Ranunculaceae family, three putative group IX AP2/ERF genes form a distinct gene cluster. Similarly, in *E. californica*, group IX AP2/ERF genes form a few small clusters (e.g., *EcAP2/ERF2–4* and *EcAP2/ERF12/17*) (Yamada et al. 2020). Although the detailed genomic composition of the group IX AP2/ERF gene cluster in *Coptis* species remains to be fully elucidated, our findings suggest that group IX AP2/ERF gene clusters are conserved in basal eudicots. Furthermore, these findings raise the possibility that AP2/ERF gene clusters have been independently formed, and that gene duplication and functional diversification within these clusters may contribute to the evolution of the regulatory mechanisms involved in BIA biosynthesis.

### CjBJE transcription factors and CjbHLH1 form regulatory networks in BIA Biosynthesis

When we examined the effect of RNA silencing of each *CjBJE* gene, only *CjBJE3*-RNAi significantly reduced the expression of genes involved in berberine biosynthesis (Figure 2). In contrast, the overexpression of *CjBJE3* and *CjBJE5* resulted in a significant increase in the expression of berberine biosynthetic enzyme genes. The weak effect of *CjBJE5*-RNAi on berberine biosynthesis might have been due to the relatively low expression of *CjBJE5* in *C. japonica* 156-S cells (Figure S4). Moreover, the expression of *CjbHLH1* and *CjWRKY1*, which have been identified as positive regulators of berberine biosynthesis, was slightly reduced in *CjBJE3*-RNAi cells, possibly resulting in different effects of RNAi on gene expression levels (Figure 3A).

In addition to the reduction in the expression levels of other *CjBJE* genes by the suppression of *CjBJE3*, the overexpression of *CjBJE3* increased the expression of other *CjBJE* genes, except for *CjBJE1*. In contrast, the overexpression of *CjBJE5* unexpectedly decreased the expression of other *CjBJE* genes (Figure 4). These results indicate the presence of a regulatory loop among group IX AP2/ERF genes and the possible presence of a negative feedback mechanism. In MIA biosynthesis, the expression of clustered *ORCA* genes is regulated by other ORCA proteins. For instance, ORCA5 and ORCA6 have been reported to upregulate the expression of other *ORCA* genes (Paul et al. 2020; Singh et al. 2020). Such regulatory loops among the clustered group IX AP2/ERFs likely ensure the spatiotemporal regulation of complex specialized metabolism. Although a self-regulatory loop has been proposed for highly clustered *ORCA* genes, a similar regulatory network appears to be present among *CjBJE* genes, although they do not form distinct gene clusters such as those seen in *Coptis*. This suggests a conserved regulatory mechanism for group IX ERFs across different plant species, even in the absence of tight clusters.

Regarding the putative *cis*-elements involved in these regulatory loops, our LUC reporter assays indicated the importance of a GCC-like sequence (e.g., AGCCACC) in transactivation by CjBJE3 and CjBJE5 (Figure 5B). Although direct binding to the exact sequence remains to be fully verified using *in vitro* binding assays, several findings suggest that CjBJE3 and CjBJE5 may recognize a variety of GC-rich sequences. Indeed, our promoter analysis of *C. chinensis* revealed that, while the GCCACC sequence is not strictly conserved across the promoter regions of genes upregulated by CjBJE3 and CjBJE5, a canonical GCC-box (GCCGCC) is present within the promoters of *CjBJE2*, *CjBJE4*, and *CjBJE5* homologous genes (Figure S12). Furthermore, our promoter analysis in *E. californica* indicated that various GC-containing sequence variants were enriched among the upregulated gene promoters (Figure S11). Since the divergence of promoter sequences between *C. japonica* and *C. chinensis CAS* and *4’OMT* genes is observed (Figure S13), obtaining the genome information of *C. japonica* will be essential in future studies to identify the precise *cis*-elements and elucidate the *cis*-mediated regulatory mechanism of biosynthetic genes in this species.

Our transient analysis suggests that CjbHLH1 controls the expression of *CjBJE* genes, particularly *CjBJE3*, which may subsequently regulate other *CjBJE* family members (Figure 6). In nicotine and MIA biosynthesis, tobacco and *C. roseus* MYC2-type bHLHs NtMYC2b and CrMYC2 directly or indirectly regulate the expression of *NtERF189* and *ORCA3*, respectively (Zhang et al. 2011, 2012). Since MYC2 plays a pivotal role in JA signalling, the MeJA-induced expression of biosynthetic genes is mediated by MYC2 and MYC2-regulated group IX ERFs, although the direct binding of MYC2 to a *cis*-element in the AP2/ERF gene promoters remains unknown. The findings of the present study suggest that the regulatory networks composed of bHLH and AP2/ERF TFs have diversified during evolution, leading to the establishment of lineage-specific systems for the regulation of secondary metabolism. Whether CjbHLH1 regulates *CjBJE3* directly or indirectly remains to be confirmed in future studies. Furthermore, upregulation of *CjbHLH1* in *CjBJE3*-overexpressing protoplasts suggests the existence of a mutual regulatory loop between CjbHLH1 and CjBJE3.

### Functional conservation of BIA regulatory networks and potential applications for the exploration of uncharacterized enzymes

The positive effects of *CjBJE3* and *CjBJE5* on BIA production were confirmed in transgenic cultured *E. californica* cells. In these heterologous expression lines, the expression of numerous endogenous biosynthetic enzyme genes was upregulated (Figure 7B). Consequently, BIA end products, including chelilutine, chelirubine, macarpine, chelerythrine, and 10-hydroxychelerythrine (10-HC), were highly accumulated in both *CjBJE3*-OX and *CjBJE5*-OX cells (Figure 7C). Notably, a decrease in the levels of intermediates such as protopine and allocryptopine suggests that *CjBJE3* and *CjBJE5* effectively enhance the metabolic flow toward the end products. This accelerated metabolic flux may be further supported by the increased expression of the CYP genes (*EcMSH*, *EcP6H*, *EcDS10H*, *EcEB10H*) and *EcCPR*. Since CPR is an essential electron donor for cytochrome P450 enzymes (Rosco et al. 1997), which catalyze multiple key steps in the downstream BIA pathways, its upregulation likely facilitated the biosynthesis of end products in OX cells. Furthermore, similar to transient overexpression in *C. japonica* protoplasts, overexpression of *CjBJE3* slightly upregulated *EcbHLH1-2* expression in transgenic cultured *E. californica* cells. No clear effect was observed for *EcbHLH1-1*, consistent with previous reports showing that *EcbHLH1-2* functions as the predominant bHLH TF in *E. californica* roots and cultured cells. These results suggest that the mutual regulatory loop between BJE and bHLH is functionally conserved even in heterologous expression systems.

In both *CjBJE3*-OX and *CjBJE5*-OX cells, significant accumulation of BIA end-products was accompanied by the upregulation of numerous biosynthetic enzyme genes. Heat map analysis confirmed that many genes encoding CYPs (including the CYP82 family), OMTs, and NMTs showed increased expression in the OX lines (Figure 8). Further characterization revealed that GC-containing sequences were distributed in the promoter regions of genes that were upregulated in both *CjBJE3*-OX and *CjBJE5*-OX cells. Recent high-quality genome analysis of *E. californica* revealed that the tandem duplication of biosynthetic enzyme genes contributes to the metabolic diversity of BIAs (Rössner et al. 2026). The coordinated regulation of *CYP82* genes by CjBJE3 or CjBJE5 suggests that the regulatory system may be conserved among the clustered genes. Furthermore, clustered genes such as *DB10H_L*, *DB10H_L2*, and *CYP82P5* within the *CYP82* gene cluster were not only markedly upregulated in OX cell lines and harbored conserved GC-rich motifs in their promoter regions (Figure S11). These findings suggest that CYP enzymes are directly involved in the biosynthesis of benzophenanthridine-type BIAs. In addition, these results indicate that cells in which metabolic profiles are altered by the heterologous expression of *CjBJE* genes serve as a valuable resource for exploring novel biosynthetic enzymes. However, the catalytic activities of these candidate enzymes have not yet been fully characterized. Whereas biochemical validation using a heterologous expression system is currently ongoing, obtaining specific BIA intermediates for use as substrates remains a critical step toward elucidating their enzymatic functions.

While group IX AP2/ERF TFs involved in the regulation of specialized metabolism have been widely reported, the regulatory networks, including group IX AP2/ERFs and other TFs, are highly diversified. Although the MYC2-type bHLH TFs and group IXa AP2/ERF TFs predominantly function as core regulators of other alkaloid biosynthetic pathways, *C. japonica* employs a distinct regulatory system. In the present study, CjbHLH1, a unique, non-MYC2-type bHLH TF found in BIA-producing plant species, operated in concert with group IXb and IXc AP2/ERF TFs, represented by CjBJE3 and CjBJE5, respectively. In particular, CjBJE3, a primary regulator in *C. japonica*, orchestrates the expression of other *CjBJE* genes, and its expression is regulated by CjbHLH1. Our findings indicate that BIA-producing species have developed lineage-specific regulatory systems during evolution compared to the biosynthesis of other alkaloids (Figure S14). Future investigations to elucidate the genomic architecture of the *BJE* cluster in *C. japonica*, as well as the relationship between CjBJEs, CjbHLH1, and the core JA-signalling complex (COI1-JAZ-MYC2), will provide deeper insights into the diversification of regulatory mechanisms in BIA biosynthesis.

## Materials and methods

### Plant materials

High-berberine-producing *C. japonica* (156-S) and low-berberine-producing *C. japonica* (CjY) cells were maintained as described previously (Yamada and Sato 2016). 156-S cells for the analysis of MeJA responsiveness were grown for 2–3 weeks, and then MeJA (final concentration: 100 µM) or the same volume of dimethyl sulfoxide (DMSO) was added. After 0, 6, and 24 h culture, the cells were harvested and stored at −80□. For tissue expression analysis, three *C. japonica* Makino var. dissecta plants (from Sannan-cho, Tamba City and grown in the Medicinal Botanical Garden of Kobe Pharmaceutical University) were separated into leaf blades, petioles, roots, and rhizomes, and these parts were stored at −80□.

### Isolation of BJE genes from C. japonica

Degenerate PCR was carried out with primers based on the conserved amino acid sequences of group IX AP2/ERF proteins in *E. californica* and other Ranunculaceae plants using template cDNAs synthesized from total RNA of cultured *C. japonica* cells treated with 100 µM MeJA. Sequences of the degenerate primers are listed in Supplementary Table S1. Five full-length sequences including 5’ and 3’ UTRs were independently isolated using GeneRacer Kit with SuperScript III RT (Thermo Fisher Scientific, Waltham, MA, USA) and gene-specific primers. Sequence information has been deposited in the GenBank/DDBJ repository with accession numbers LC941495 (*CjBJE1*), LC941496 (*CjBJE2*), LC941497 (*CjBJE3*), LC941498 (*CjBJE4*), and LC941499 (*CjBJE5*).

### Vector construction

The 35S:*BJE1-sGFP*, 35S:*BJE2-sGFP*, 35S:*BJE3-sGFP* and 35S:*BJE5-sGFP* vectors were constructed to introduce each *Sal*I/*Nco*I fragment of *BJE* without a stop codon into the pUC18-*sGFP* vector, whereas the 35S:*BJE4*-sGFP vector was created to insert *sGFP* gene into 35S:*BJE4* without a stop codon using a *Sac*I site. Transient overexpression vectors (35S::*BJE3* and 35S::*BJE5*) were constructed from pBI221 by replacing the *GUS* gene with *Bam*HI (*BJE3* and *BJE5*) and *Sac*I fused full-length cDNAs of *BJE* genes. The 35S::*BJE3-SRDX* and 35S::*GIXE5-SRDX* vectors were generated to introduce both strands of DNA fragments corresponding to SRDX (LDLDLELRLGFA) with a stop codon into the overexpression vectors after removing the stop codon of *BJE* gene by inverse PCR. Expression vectors of GST fusion proteins were constructed to introduce truncated *CjBJE3* (4–186 bp was truncated; *CjBJE3*Δ61) and full-length *CjBJE5* into the pCold GST vector (TAKARA BIO Inc., Shiga, Japan) cut with the *Nde*I and *Sal*I sites using an In-Fusion HD Cloning kit (TAKARA BIO Inc.). Overexpression vectors for generating stable transformants were constructed as follows: 35S::*BJE3*, 35S::*BJE5*, 35S::*BJE3-SRDX*, and 35S::*BJE5-SRDX* vectors were digested with *Bam*HI and *Sac*I, and the resulting fragments were inserted into a pBIE binary vector digested with the same restriction enzymes.

### Phylogenetic analysis

Full-length amino acid sequences of group IX AP2/ERF proteins were retrieved from NCBI (https://www.ncbi.nlm.nih.gov/), and their conserved AP2/ERF domain regions were extracted using SMART (http://smart.embl-heidelberg.de/) (Table S2). Multiple sequence alignments of domain sequences were performed using MAFFT ver. 7 (https://mafft.cbrc.jp/alignment/server/index.html) using the default parameters (Katoh et al. 2019). A maximum-likelihood (ML) phylogenetic tree was constructed using MEGA12 (Kumar et al. 2024), based on the Jones-Taylor-Thornton (JTT) model, with gamma-distributed rates among sites (JTT+G, five discrete categories). Position filtering was performed using a 95% partial deletion cutoff to remove alignment gaps and missing data. Node support was evaluated using bootstrap analysis with 1,000 replicates. Alignment and tree files are provided in the Supplemental Dataset S1 and Supplemental Dataset S2, respectively.

### Quantitative RT-PCR

Total RNA was extracted using the RNeasy Plant Mini Kit (Qiagen, Hilden, Germany). For transient assays in 156-S protoplasts and 156-S/CjY comparisons, cDNA was synthesized from 400–1000 ng of total RNA using a PrimeScript RT reagent Kit with gDNA Eraser (TAKARA BIO Inc.), and quantitative RT-PCR (qPCR) was performed using iQ SYBR Green Supermix (SsoAdvanced) and a CFX96 Real-Time PCR Detection System (Bio-Rad, Hercules, CA, USA). For MeJA-responsiveness, tissue expression, and stable transformant analyses, cDNA was synthesized using ReverTra Ace qPCR RT Master Mix (TOYOBO Co., Ltd., Osaka, Japan), and qPCR was performed using THUNDERBIRD Next SYBR qPCR Mix (TOYOBO Co., Ltd.) on a LightCycler 96 instrument (Roche, Basel, Switzerland). The primers used are listed in Supplementary Table S3. Transcript levels were normalized to β*-actin* or α*-tubulin* and calculated using the standard curve method, except for the tissue expression analysis, for which we utilized the 2^-Ct^ method. Further details followed previous reports (Yamada et al. 2020).

### Subcellular localization

The 35S::*CjBJEs-sGFP* genes were introduced into 156-S protoplasts, as previously described (Yamada and Sato 2016). The localization of the sGFP fusion proteins was detected using a BZ-9000 fluorescence microscope (KEYENCE, Osaka, Japan).

### Transient RNAi and overexpression in C. japonica protoplasts

The target regions for RNAi-mediated suppression of *CjBJE* genes are shown in Supplementary Figure S14, whereas the target for *CjbHLH1* was described previously (Yamada et al. 2011a). In vitro dsRNA synthesis was performed using a T7 RiboMAX Express RNA Production System (Promega, Madison, WI, USA) with PCR-generated templates containing T7 promoter sequences at both ends (Table S4). Protoplasts were prepared from 2–3-week-old 156-S cultured cells, and dsRNA was introduced by polyethylene glycol (PEG)-mediated transfection according to established protocols (Dubouzet et al. 2005; Kato et al. 2007; Yamada et al. 2011a). The dsRNA of the GFP sequence was used as the control. After 72 h of incubation, total RNA was extracted for further analysis.

For transient overexpression, protoplasts were transfected with *35S::BJE3*, *35S::BJE5*, or *35S::CjbHLH1* vectors and incubated for 24 h before RNA extraction.

### Dual-Luciferase reporter assay

35S::*BJE3* and 35S::*BJE5* plasmids as effector constructs, *4’OMT* and *CHS* promoter::*PpLUC* plasmids as reporter constructs, and 35S::*RrLUC* plasmid as a reference construct into 156-S protoplasts were introduced into *C. japonica* 156-S protoplasts by PEG-mediated transfection. Transactivation activities were analyzed using a Dual-Luciferase Reporter Assay System (Promega, Madison, WI, USA) and a GloMax 20/20 Luminometer (Promega) as previously reported (Yamada et al. 2020). All experiments were performed with three biological transfections.

### Electrophoresis mobility shift assay

GST-recombinant proteins were expressed in *E. coli* BL21 cells (TAKARA BIO Inc.) by inducing with 0.1 mM isopropyl-β-D-thiogalactoside (IPTG) for 24 h at 15°C and were purified using Glutathione Sepharose 4B (Cytiva). EMSA was performed as previously described, using a LightShift Chemiluminescent EMSA Kit (Thermo Fisher Scientific) with minor modifications (Yamada et al. 2016). Biotin-labelled probes (80 fmol) and the purified recombinant proteins (2 μg) were mixed with 1× binding buffer [5 mM MgCl_2_, 50 mM KCl, 0.01% NP-40, 1.25% glycerol, 0.05 μg/μ l poly (dI-dC)] from the Kit (Thermo Fisher Scientific). After a 20 min incubation at 4°C, the reaction mixtures were separated on a 5% polyacrylamide gel and transblotted onto zeta-probe membranes (Bio-Rad) using a Trans-Blot SD Semi-Dry Electrophoretic Transfer Cell (Bio-Rad) at 380 mA for 30 min. Shifts in the biotinylated probes were detected using an ImageQuant LAS4010 CCD Imaging System (Cytiva). Nucleotide sequences of the probes used are listed in Supplementary Table S5.

### Transformation of California poppy cells

The petioles of California poppy plants (cultivar Hitoezaki, Takii Seed Co., Ltd., Kyoto, Japan) were cut into 5±10 mm segments and used for *Agrobacterium tumefaciens* (LBA4404)-mediated transformation, as previously described (Yamada et al. 2015). After approximately 6 months of selection on agar medium containing 150 μg/ml kanamycin and 200 μg/ml cefotaxime, the obtained cells were cultured in liquid Linsmaier–Skoog (LS) medium (pH5.7) containing 3% sucrose, 10 μM 1-naphthylacetic acid and 1 μM 6-benzyladenine every 2–3 weeks. Approximately 0.3 g fresh weight (FW) of California poppy cells was inoculated in 12.5 mL of LS medium and cultured for 10 days.

### RNA sequencing

Total RNA was extracted from the cultured cells using the RNeasy Plant Mini Kit and RNase-Free DNase Set (Qiagen). RNA sequencing and mapping analyses were performed by Eurofins Genomics K.K. (Tokyo, Japan) according to the following procedures. Sequencing was performed using an Illumina NovaSeq 6000 (San Diego, CA, USA) and cleaned using Trimmomatic ver. 0.39. Read mapping was performed using BWA ver. 0.7.17 and SAMtools ver. 1.10, and expression levels were quantified by the TMM method using edgeR ver. 3.16.5. The sequencing data from this study have been deposited in the DDBJ Sequence Read Archive (DRA) under the BioProject accession number PRJDB42997. Heatmaps were generated using the pheatmap package, with TMM values normalized via Z-score transformation across genes to show relative expression changes among samples.

### Metabolite analysis

Approximately 0.1 g FW of cells were ground and extracted in 4 μL/mg methanol containing 0.01 N HCl at room temperature (25°C) for 24 h, and the supernatant was collected for subsequent analysis. The culture medium samples were concentrated to 3 mL using Sep-Pak Plus C18 cartridges (Waters, Milford, MA, USA). All samples were analyzed using an ACQUITY UPLC-MS system coupled with a QDa mass detector (Waters), as described by Yamada et al. 2024, with slight modifications.

Separation was performed on an ACQUITY UPLC BEH C18 column (2.1×100 mm, 1.7 μm). For the analysis of cultured cells and medium, a gradient of water (solvent A) and acetonitrile (solvent B), both containing 0.1% formic acid, was applied: 0–1 min, 10–15% B; 1–12 min, 15–50% B; 12–13.5 min, 50–80% B; 13.5–15 min, 80–10% B; and 15–16 min, 10% B. Qda mass spectrometer settings have been described previously (Yamada et al. 2024, 2025). The chromatographic peaks were identified by comparing their retention times and *m/z* values with those of authentic standards. As authentic standards for chelirubine, macarpine, 10-HC, and chelilutine were not available, they were identified based on *m/z* and literature data (Takemura et al. 2010). Alkaloid content was calculated using the standard curves of authentic standards; specifically, chelirubine, macarpine, 10-HC, and chelilutine were quantified using the standard curves of sanguinarine and chelerythrine.

Sanguinarine chloride and magnocurcumin were purchased from Sigma-Aldrich (St. Louis, MO, USA). Chelerythrine chloride and protopine were purchased from Tokyo Chemical Industry Co. Ltd. (Tokyo, Japan). Allocyptopine was purchased from Cayman Chemical Co. (Ann Arbor, MI, USA). Cheilanthifoline was obtained from ChemFaces Biochemical Co. Ltd. (Wuhan, China). Scoulerine was obtained from Toronto Research Chemicals, Inc. (Toronto, ON, Canada). Reticuline was provided by MITSUI CHEMICALS, Inc. (Tokyo, Japan).

### Promoter analysis

Genomic DNA sequences spanning 2000 bp upstream from the 1^st^ ATG of the candidate genes were retrieved using the Biomart tool in Phytozome v14 (https://phytozome-next.jgi.doe.gov/). Motif prediction within these upstream promoter regions was performed using MEME Suite ver. 5.5.9 (https://meme-suite.org/meme/tools/meme).

## Supplementary materials

The following supplementary materials are available online.

Supplementary Figure S1. Transient RNAi screening of candidate *AP2/ERF* genes found in *C. japonica* expressed sequence tags.

Supplementary Figure S2. Amino acid sequence alignment of CjBJEs with other group IX AP2/ERF transcription factors involved in alkaloid biosynthesis regulation.

Supplementary Figure S3. Tissue-specific expression levels of *CjBJE* genes in *Coptis japonica*.

Supplementary Figure S4. Expression levels and copy numbers of *CjBJE* genes in *Coptis japonica* cultured cells.

Supplementary Figure S5. Subcellular localization of CjBJE proteins in *Coptis japonica* protoplasts.

Supplementary Figure S6. Transient RNA silencing of *CjBJE* genes in *Coptis japonica* protoplasts.

Supplementary Figure S7. Coomassie brilliant blue-stained purified GST, GST-CjBJE3, GST-CjBJE3Δ61, and GST-CjBJE5 proteins.

Supplementary Figure S8. Electrophoresis mobility shift assay using GST-fused recombinant proteins.

Supplementary Figure S9. Establishment of transgenic cultured cell lines of the California poppy.

Supplementary Figure S10. The overexpression of *CjBJE3*_SRDX and *CjBJE5*_SRDX in cultured *Eschscholzia californica* cells.

Supplementary Figure S11. MEME motif analysis of promoter regions in BIA biosynthetic genes.

Supplementary Figure S12. Nucleotide sequences of *Coptis chinensis BJE2*, *BJE4*, and *BJE5* homologous genes.

Supplementary Figure S13. Nucleotide sequence alignment of *Coptis japonica* and *Coptis chinensis 4’OMT* and *CAS* promoter regions.

Supplementary Figure S14. Proposed model for the transcriptional regulation of BIA biosynthesis mediated by CjbHLH1 and CjBJE transcription factors in *Coptis japonica*.

Supplementary Figure S15. Target regions for transient RNAi in each *CjBJE* gene.

Supplemental Dataset S1. Multiple sequence alignment used for phylogenetic analysis of AP2/ERF domains (FASTA format).

Supplemental Dataset S2. Phylogenetic tree file of group IX AP2/ERF transcription factors (Newick format).

Supplemental Dataset S3. RNA-seq expression values (TMM) for selected BIA biosynthesis and transcription factor genes (Excel).

Supplemental Dataset S4. Quantification of benzylisoquinoline alkaloids in transgenic California poppy cultured cells (Excel format).

Supplemental Dataset S5. UPLC-MS chromatograms of cell extracts and culture media (PDF format).

## Acknowledgements

We thank Ms. Yoko Nakahara, Ms. Mari Hiratani, Mr. Yuto Hatta, and Mr. Yuki Mori (Kobe Pharmaceutical University, Japan) for their assistance with preliminary experiments. We also thank Ms. Azusa Hirano, Mr. Takahiro Ohi, and Dr. Yumi Nishiyama (Medicinal Botanical Garden, Kobe Pharmaceutical University, Japan) for their support in growing the *C. japonica* plants. We are grateful to the Hyogo Prefectural Technology Center for Agriculture, Forestry, and Fisheries for kindly providing the *C. japonica* var dissecta plants. In addition, we thank Dr. Kazufumi Yazaki (Kyoto University, Japan) for providing the low-berberine-producing cultured *C. japonica* cells, and Dr. Yoshito Ikeda (Shiga University of Medical Science, Japan) for fruitful discussions regarding this study. We would like to thank Editage (www.editage.jp) for English language editing.

## Author contributions

**Y.Y.:** Conceptualization, isolation of *BJE* genes, vector construction, gene expression analysis, dual-LUC reporter assays in *C. japonica*, EMSA, generation of *E. californica* transformants, transcriptome and metabolite analyses of transgenic *E. californica* cells, data curation, and writing and editing of the manuscript. **Y.T.:** Support for the GST fusion protein expression and EMSA. **A.I.:** Support for the qRT-PCR analysis. **N.S.:** Conceptualization, and reviewing and editing of the manuscript. **F.S.:** Conceptualization, reviewing and editing of the manuscript, and project administration. All the authors have read and approved the final version of the manuscript.

## Data availability

The raw RNA-sequencing data generated in this study have been deposited in the DDBJ Sequence Read Archive (DRA) under BioProject ID PRJDB42997 (DRR Run: DRR1089032-DRR1089044). All other data underlying this article are available in the article and in its online supplementary material.

## References

Apuya NR, Park J-H, Zhang L, Ahyow M, Davidow P, Van Fleet J, Rarang JC, Hippley M, Johnson TW, Yoo H-D, et al. Enhancement of alkaloid production in opium and California poppy by transactivation using heterologous regulatory factors. Plant Biotechnol J. 2008:6(2):160–175.

Becker A, Yamada Y, and Sato F. California poppy (Eschscholzia californica), the Papaveraceae golden girl model organism for evodevo and specialized metabolism. Front Plant Sci. 2023:14:1084358.

Cárdenas PD, Sonawane PD, Pollier J, Vanden Bossche R, Dewangan V, Weithorn E, Tal L, Meir S, Rogachev I, Malitsky S, et al. GAME9 regulates the biosynthesis of steroidal alkaloids and upstream isoprenoids in the plant mevalonate pathway. Nat Commun. 2016:7:10654.

Chen D-X, Pan Y, Wang Y, Cui Y-Z, Zhang Y-J, Mo R-Y, Wu X-L, Tan J, Zhang J, Guo L-A, et al. The chromosome-level reference genome of Coptis chinensis provides insights into genomic evolution and berberine biosynthesis. Hortic Res. 2021:8(1):121.

De Boer K, Tilleman S, Pauwels L, Vanden Bossche R, De Sutter V, Vanderhaeghen R, Hilson P, Hamill JD, and Goossens A. APETALA2/ETHYLENE RESPONSE FACTOR and basic helix-loop-helix tobacco transcription factors cooperatively mediate jasmonate-elicited nicotine biosynthesis. Plant J. 2011:66(6):1053–1065.

De Geyter N, Gholami A, Goormachtig S, and Goossens A. Transcriptional machineries in jasmonate-elicited plant secondary metabolism. Trends Plant Sci. 2012:17(6):349–359.

Dubouzet JG, Morishige T, Fujii N, An C-I, Fukusaki E-I, Ifuku K, and Sato F. Transient RNA silencing of scoulerine 9-O-methyltransferase expression by double stranded RNA in Coptis japonica protoplasts. Biosci Biotechnol Biochem. 2005:69(1):63–70.

Facchini PJ. ALKALOID BIOSYNTHESIS IN PLANTS: Biochemistry, cell biology, molecular regulation, and metabolic engineering applications. Annu Rev Plant Physiol Plant Mol Biol. 2001:52(1):29–66.

van der Fits L and Memelink J. ORCA3, a jasmonate-responsive transcriptional regulator of plant primary and secondary metabolism. Science. 2000:289(5477):295–297.

He S-M, Liang Y-L, Cong K, Chen G, Zhao X, Zhao Q-M, Zhang J-J, Wang X, Dong Y, Yang J-L, et al. Identification and characterization of genes involved in benzylisoquinoline alkaloid biosynthesis in Coptis species. Front Plant Sci. 2018:9:731.

Hiratsu K, Matsui K, Koyama T, and Ohme-Takagi M. Dominant repression of target genes by chimeric repressors that include the EAR motif, a repression domain, in Arabidopsis: Dominant repression by EAR-motif chimeric repressor. Plant J. 2003:34(5):733–739.

Hori K, Yamada Y, Purwanto R, Minakuchi Y, Toyoda A, Hirakawa H, and Sato F. Mining of the Uncharacterized Cytochrome P450 Genes Involved in Alkaloid Biosynthesis in California Poppy Using a Draft Genome Sequence. Plant Cell Physiol. 2018:59(2):222–233.

Inui T, Kawano N, Shitan N, Yazaki K, Kiuchi F, Kawahara N, Sato F, and Yoshimatsu K. Improvement of benzylisoquinoline alkaloid productivity by overexpression of 3’-hydroxy-N-methylcoclaurine 4’-O-methyltransferase in transgenic Coptis japonica plants. Biol Pharm Bull. 2012:35(5):650–659.

Kato N, Dubouzet E, Kokabu Y, Yoshida S, Taniguchi Y, Dubouzet JG, Yazaki K, and Sato F. Identification of a WRKY protein as a transcriptional regulator of benzylisoquinoline alkaloid biosynthesis in Coptis japonica. Plant Cell Physiol. 2007:48(1):8–18.

Katoh K, Rozewicki J, and Yamada KD. MAFFT online service: multiple sequence alignment, interactive sequence choice and visualization. Brief Bioinform. 2019:20(4):1160–1166.

Kumar S, Stecher G, Suleski M, Sanderford M, Sharma S, and Tamura K. MEGA12: Molecular evolutionary genetic analysis version 12 for adaptive and green computing. Mol Biol Evol. 2024:41(12):msae263.

Licausi F, Ohme-Takagi M, and Perata P. APETALA2/Ethylene Responsive Factor (AP2/ERF) transcription factors: mediators of stress responses and developmental programs. New Phytol. 2013:199(3):639–649.

Liu W, Tian X, Feng Y, Hu J, Wang B, Chen S, Liu D, and Liu Y. Genome-wide analysis of bHLH gene family in Coptis chinensis provides insights into the regulatory role in benzylisoquinoline alkaloid biosynthesis. Plant Physiol Biochem. 2023:201(107846):107846.

Liu Y, Wang B, Shu S, Li Z, Song C, Liu D, Niu Y, Liu J, Zhang J, Liu H, et al. Analysis of the Coptis chinensis genome reveals the diversification of protoberberine-type alkaloids. Nat Commun. 2021:12(1):3276.

Lu X, Zhang L, Zhang F, Jiang W, Shen Q, Zhang L, Lv Z, Wang G, and Tang K. AaORA, a trichome-specific AP2/ERF transcription factor of *Artemisia annua*, is a positive regulator in the artemisinin biosynthetic pathway and in disease resistance to Botrytis cinerea. New Phytol. 2013:198(4):1191–1202.

Menke FL, Champion A, Kijne JW, and Memelink J. A novel jasmonate- and elicitor-responsive element in the periwinkle secondary metabolite biosynthetic gene *Str* interacts with a jasmonate- and elicitor-inducible AP2-domain transcription factor, ORCA2. EMBO J. 1999:18(16):4455–4463.

Mishra S, Triptahi V, Singh S, Phukan UJ, Gupta MM, Shanker K, and Shukla RK. Wound induced tanscriptional regulation of benzylisoquinoline pathway and characterization of wound inducible PsWRKY transcription factor from *Papaver somniferum*. PLoS One. 2013:8(1):e52784.

Mizoi J, Shinozaki K, and Yamaguchi-Shinozaki K. AP2/ERF family transcription factors in plant abiotic stress responses. Biochim Biophys Acta. 2012:1819(2):86–96.

Nakano T, Suzuki K, Fujimura T, and Shinshi H. Genome-wide analysis of the *ERF* gene family in Arabidopsis and rice. Plant Physiol. 2006:140(2):411–432.

Nakayasu M, Shioya N, Shikata M, Thagun C, Abdelkareem A, Okabe Y, Ariizumi T, Arimura G-I, Mizutani M, Ezura H, et al. JRE4 is a master transcriptional regulator of defense-related steroidal glycoalkaloids in tomato. Plant J. 2018:94(6):975–990.

Pan Q, Wang C, Xiong Z, Wang H, Fu X, Shen Q, Peng B, Ma Y, Sun X, and Tang K. CrERF5, an AP2/ERF Transcription Factor, Positively Regulates the Biosynthesis of Bisindole Alkaloids and Their Precursors in *Catharanthus roseus*. Front Plant Sci. 2019:10:931.

Paul P, Singh SK, Patra B, Liu X, Pattanaik S, and Yuan L. Mutually Regulated AP2/ERF Gene Clusters Modulate Biosynthesis of Specialized Metabolites in Plants. Plant Physiol. 2020:182(2):840–856.

Paul P, Singh SK, Patra B, Sui X, Pattanaik S, and Yuan L. A differentially regulated AP2/ERF transcription factor gene cluster acts downstream of a MAP kinase cascade to modulate terpenoid indole alkaloid biosynthesis in *Catharanthus roseus*. New Phytol. 2017:213(3):1107–1123.

Rosco A, Pauli HH, Priesner W, and Kutchan TM. Cloning and heterologous expression of NADPH-cytochrome P450 reductases from the Papaveraceae. Arch Biochem Biophys. 1997:348(2):369–377.

Rössner L-H, Rössner C, Kong D, Lotz D, Weisert A, Yamada Y, Sato F, Davies K, Rupp O, Fuchs J, et al. Gene and genome duplications have contrasting impacts on biosynthetic and flower developmental pathways in California poppy. Plant Cell. 2026:38(3):koag039.

Sato F. Characterization of plant functions using cultured plant cells, and biotechnological applications. Biosci Biotechnol Biochem. 2013:77(1):1–9.

Sato F. Plant Alkaloid Engineering.. In. Comprehensive Natural Products III, H-W (ben) Liu and TP Begley, eds. (Elsevier: Oxford), pp. 700–755.

Shoji T, Kajikawa M, and Hashimoto T. Clustered transcription factor genes regulate nicotine biosynthesis in tobacco. Plant Cell. 2010:22(10):3390–3409.

Shoji T and Yuan L. ERF Gene Clusters: Working Together to Regulate Metabolism. Trends Plant Sci. 2021:26(1):23–32.

Singh A, Menéndez-Perdomo IM, and Facchini PJ. Benzylisoquinoline alkaloid biosynthesis in opium poppy: an update. Phytochem Rev. 2019:18(6):1457–1482.

Singh SK, Patra B, Paul P, Liu Y, Pattanaik S, and Yuan L. Revisiting the ORCA gene cluster that regulates terpenoid indole alkaloid biosynthesis in *Catharanthus roseus*. Plant Sci. 2020:293:110408.

Takemura T, Ikezawa N, Iwasa K, and Sato F. Metabolic diversification of benzylisoquinoline alkaloid biosynthesis through the introduction of a branch pathway in *Eschscholzia californica*. Plant Cell Physiol. 2010:51(6):949–959.

Thagun C, Imanishi S, Kudo T, Nakabayashi R, Ohyama K, Mori T, Kawamoto K, Nakamura Y, Katayama M, Nonaka S, et al. Jasmonate-Responsive ERF Transcription Factors Regulate Steroidal Glycoalkaloid Biosynthesis in Tomato. Plant Cell Physiol. 2016:57(5):961–975.

Udomsom N, Rai A, Suzuki H, Okuyama J, Imai R, Mori T, Nakabayashi R, Saito K, and Yamazaki M. Function of AP2/ERF Transcription Factors Involved in the Regulation of Specialized Metabolism in *Ophiorrhiza pumila* Revealed by Transcriptomics and Metabolomics. Front Plant Sci. 2016:7:1861.

Yamada Y, Hirakawa H, Hori K, Minakuchi Y, Toyoda A, Shitan N, and Sato F. Comparative analysis using the draft genome sequence of California poppy (*Eschscholzia californica*) for exploring the candidate genes involved in benzylisoquinoline alkaloid biosynthesis. Biosci Biotechnol Biochem. 2021:85(4):851–859.

Yamada Y, Kato N, Kokabu Y, Luo Q, Dubouzet JG, and Sato F. Identification of regulatory protein genes involved in alkaloid biosynthesis using a transient RNAi system. Methods Mol Biol. 2010:643:33–45.

Yamada Y, Kokabu Y, Chaki K, Yoshimoto T, Ohgaki M, Yoshida S, Kato N, Koyama T, and Sato F. Isoquinoline alkaloid biosynthesis is regulated by a unique bHLH-type transcription factor in *Coptis japonica*. Plant Cell Physiol. 2011a:52(7):1131–1141.

Yamada Y, Koyama T, and Sato F. Basic helix-loop-helix transcription factors and regulation of alkaloid biosynthesis. Plant Signal Behav. 2011b:6(11):1627–1630.

Yamada Y, Matsui T, Nomura F, Shimizu Y, Oguni T, Murakami Y, Shitan N, Terasaka K, and Sato F. Molecular characterization of two *O*-methyltransferases involved in benzylisoquinoline alkaloid biosynthesis in *Aristolochia debilis*. Plant Physiol Biochem. 2025:230(110956):110956.

Yamada Y, Motomura Y, and Sato F. CjbHLH1 homologs regulate sanguinarine biosynthesis in *Eschscholzia californica* cells. Plant Cell Physiol. 2015:56(5):1019–1030.

Yamada Y, Nishida S, Shitan N, and Sato F. Genome-wide identification of AP2/ERF transcription factor-encoding genes in California poppy (*Eschscholzia californica*) and their expression profiles in response to methyl jasmonate. Sci Rep. 2020:10(1):18066.

Yamada Y and Sato F. Transcription factors in alkaloid biosynthesis. Int Rev Cell Mol Biol. 2013:305:339–382.

Yamada Y and Sato F. Tyrosine phosphorylation and protein degradation control the transcriptional activity of WRKY involved in benzylisoquinoline alkaloid biosynthesis. Sci Rep. 2016:6:31988.

Yamada Y and Sato F. Transcription factors in alkaloid engineering. Biomolecules. 2021:11(11):1719.

Yamada Y, Shimada T, Motomura Y, and Sato F. Modulation of benzylisoquinoline alkaloid biosynthesis by heterologous expression of CjWRKY1 in *Eschscholzia californica* cells. PLoS One. 2017:12(10):e0186953.

Yamada Y, Tamagaki E, Shitan N, and Sato F. Integrated metabolite profiling and transcriptome analysis reveal candidate genes involved in the biosynthesis of benzylisoquinoline alkaloids in *Corydalis solida*. Plant Biotechnol (Tsukuba). 2024:41:267–276.

Yamada Y, Yoshimoto T, Yoshida ST, and Sato F. Characterization of the Promoter Region of Biosynthetic Enzyme Genes Involved in Berberine Biosynthesis in *Coptis japonica*. Front Plant Sci. 2016:7:1352.

Yu Z-X, Li J-X, Yang C-Q, Hu W-L, Wang L-J, and Chen X-Y. The jasmonate-responsive AP2/ERF transcription factors AaERF1 and AaERF2 positively regulate artemisinin biosynthesis in *Artemisia annua* L. Mol Plant. 2012:5(2):353–365.

Zhang H, Hedhili S, Montiel G, Zhang Y, Chatel G, Pré M, Gantet P, and Memelink J. The basic helix-loop-helix transcription factor CrMYC2 controls the jasmonate-responsive expression of the ORCA genes that regulate alkaloid biosynthesis in *Catharanthus roseus*. Plant J. 2011:67(1):61–71.

Zhang H-B, Bokowiec MT, Rushton PJ, Han S-C, and Timko MP. Tobacco transcription factors NtMYC2a and NtMYC2b form nuclear complexes with the NtJAZ1 repressor and regulate multiple jasmonate-inducible steps in nicotine biosynthesis. Mol Plant. 2012:5(1):73–84.

Zhang M, Lu P, Zheng Y, Huang X, Liu J, Yan H, Quan H, Tan R, Ren F, Jiang H, et al. Genome-wide identification of AP2/ERF gene family in *Coptis Chinensis* Franch reveals its role in tissue-specific accumulation of benzylisoquinoline alkaloids. BMC Genomics. 2024:25(1):972.

