## Supplementary materials for "Jasmonate-responsive group IX AP2/ERF transcription factors control the biosynthesis of benzylisoquinoline alkaloids"

**Corresponding Authors:** Yasuyuki Yamada*

**Corresponding Authors:** Fumihiko Sato*

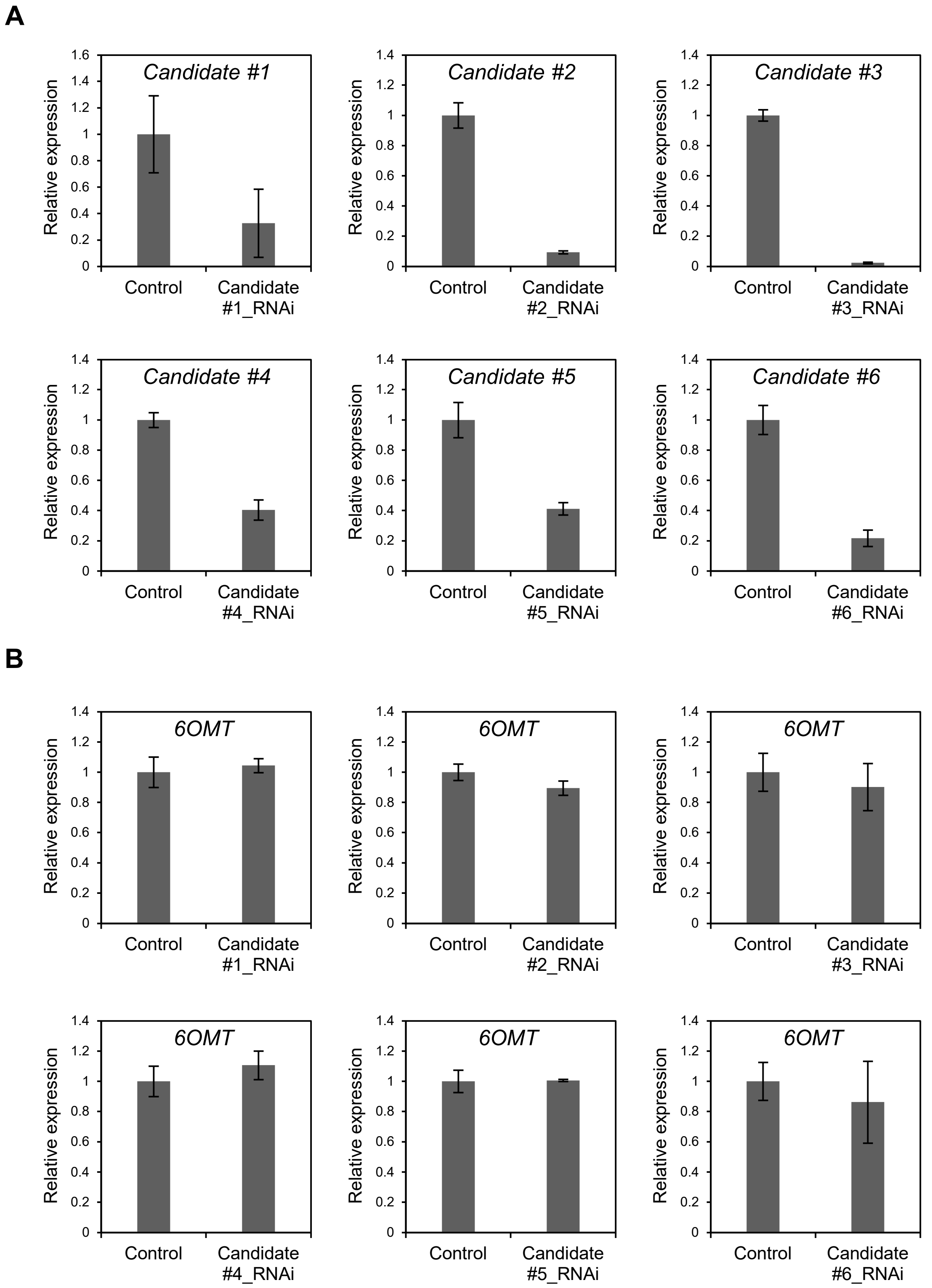


**Supplementary Figure S1. Transient RNAi screening of candidate *AP2/ERF* genes found in *C. japonica* expressed sequence tags.**
Relative transcript levels of six candidate *AP2/ERF* genes (A) and *Cj6OMT* (B) were determined by qRT-PCR and normalized to the *CjATPase* gene. Relative expression levels are shown in terms of fold-change compared with control (set to 1.0). Error bars indicate standard deviation of three biological replicates.

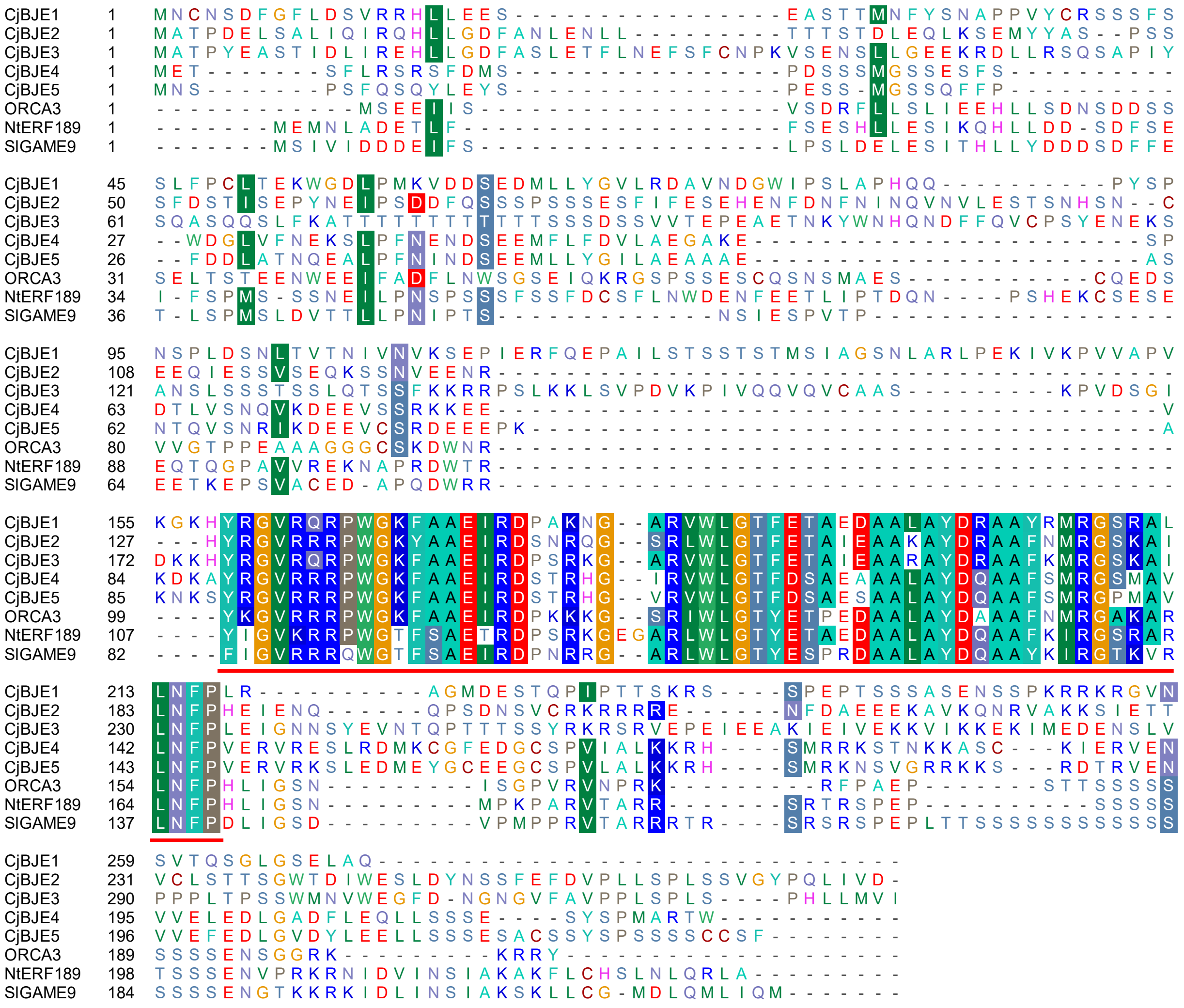

**Supplementary Figure S2. Amino acid sequence alignment of CjBJEs with other group IX AP2/ERF transcription factors involved in alkaloid biosynthesis regulation.**

Sequences were aligned using BioEdit. Conserved AP2/ERF domain is indicated in red.

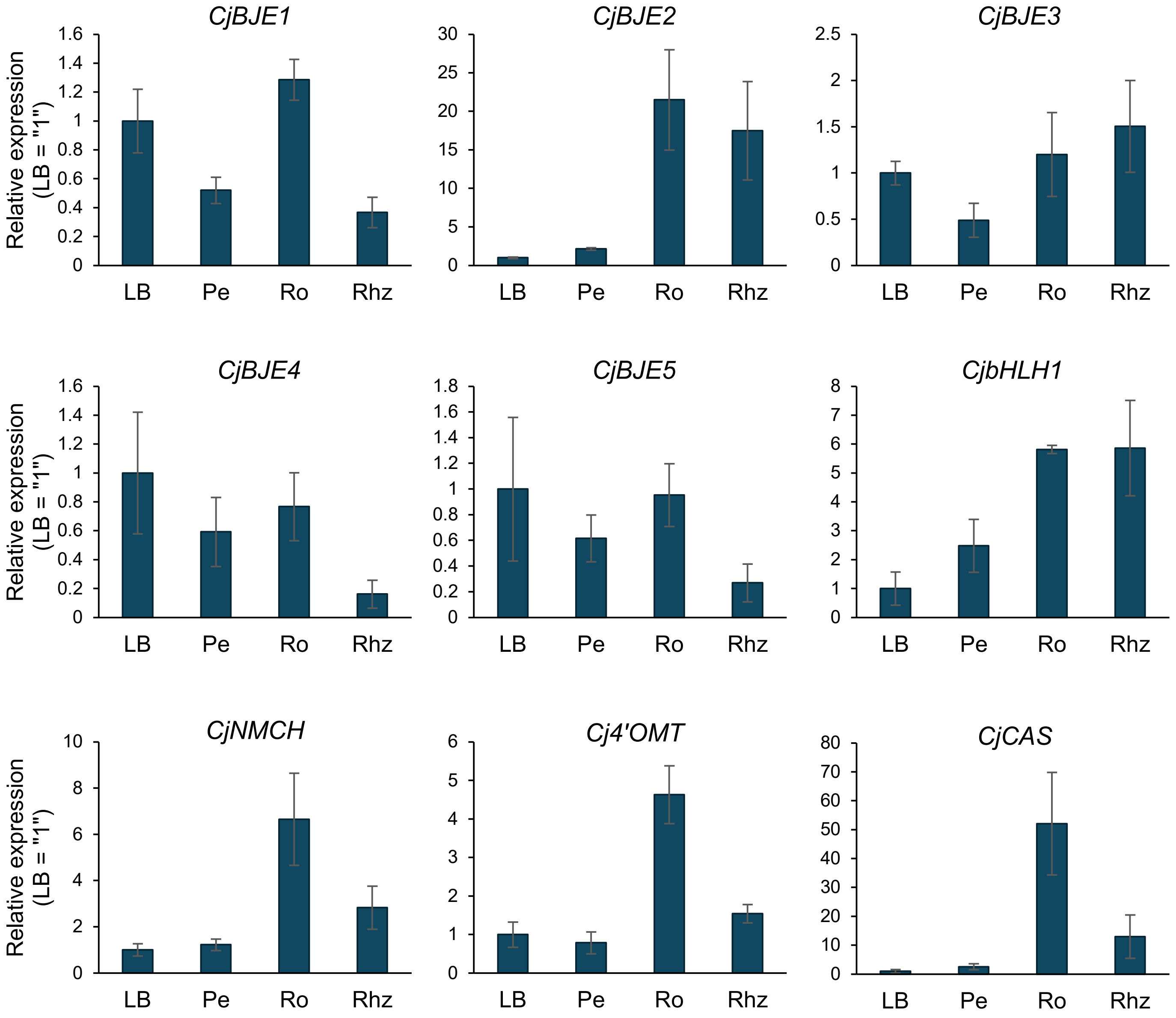


**Supplementary Figure S3. Tissue-specific expression levels of *CjBJE* genes in *Coptis japonica*.**
Expression levels of *CjBJE1–5*, *CjbHLH1*, and other biosynthetic enzyme genes (*NMCH*, *4'OMT*, and *CAS*) were determined by qRT-PCR. Relative transcript levels were calculated using *CjACTIN* as an internal control and are shown in terms of fold-changes compared to leaf blades (set to 1.0). Error bars indicate standard deviation of three biological replicates. LB, leaf blades; Pe, petiole; Ro, root; Rhz, rhizome.

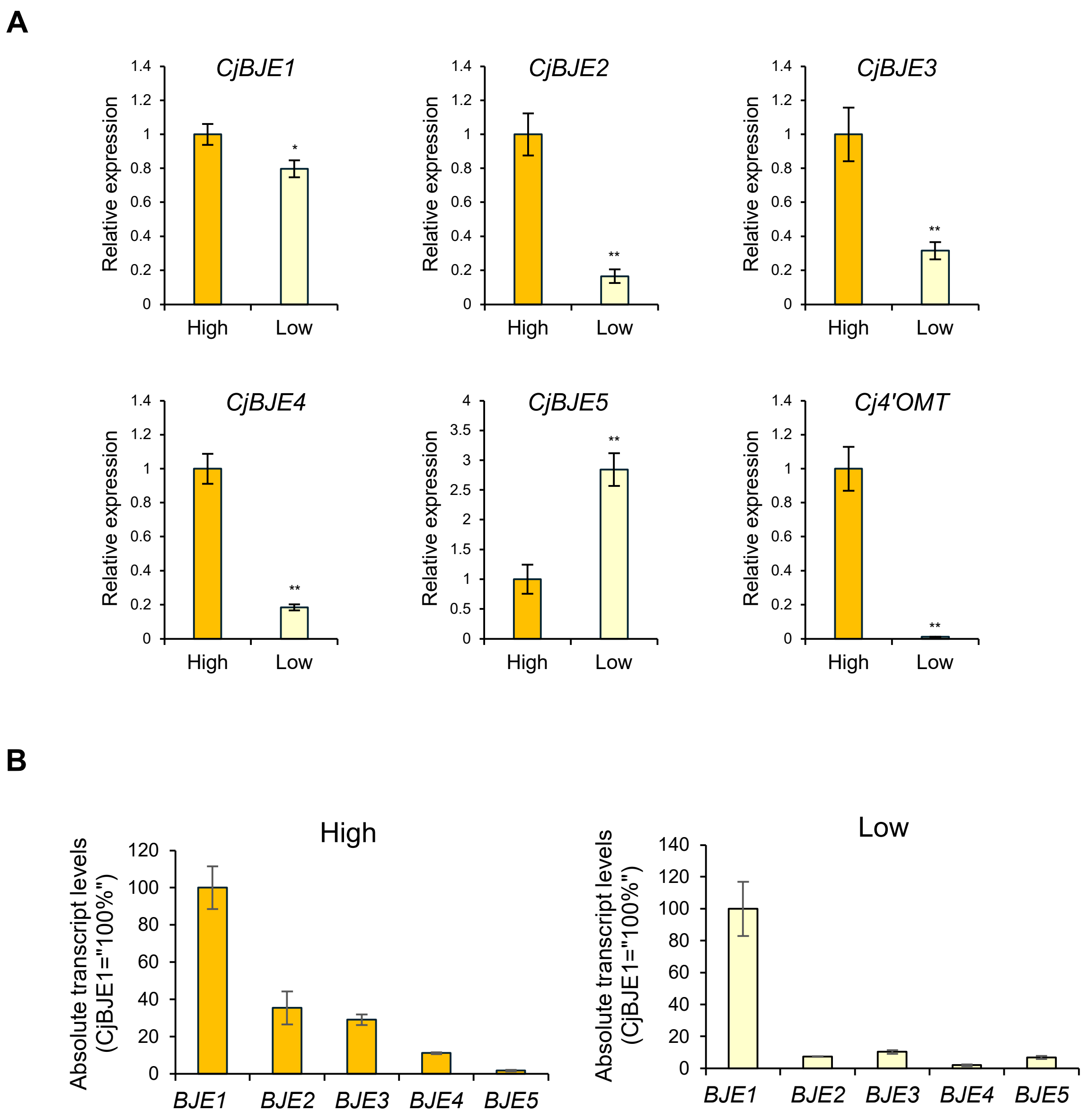


**Supplementary Figure S4. Expression levels and copy numbers of *CjBJE* genes in *Coptis japonica* cultured cells.**

(A) Expression levels of *CjBJE1–5* and *Cj4'OMT* in berberine high-producing cells (High) and low-producing cells (Low) were determined by qRT-PCR. Relative transcript levels were calculated using *CjACTIN* as an internal control and are shown in terms of fold-change compared with high-producing cells (set to 1.0). Error bars indicate standard deviations from three biological replicates. Asterisks indicate significant differences compared with control (Student's *t*-test: *$p < 0.05$, **$p < 0.01$). (B) Absolute quantification of copy numbers for *CjBJE1–5* in both cultured cells. Copy numbers were determined by real-time PCR using plasmid DNA standard curves. Data are represented as relative copy numbers normalized to that of *CjBJE1* (set to 100%).

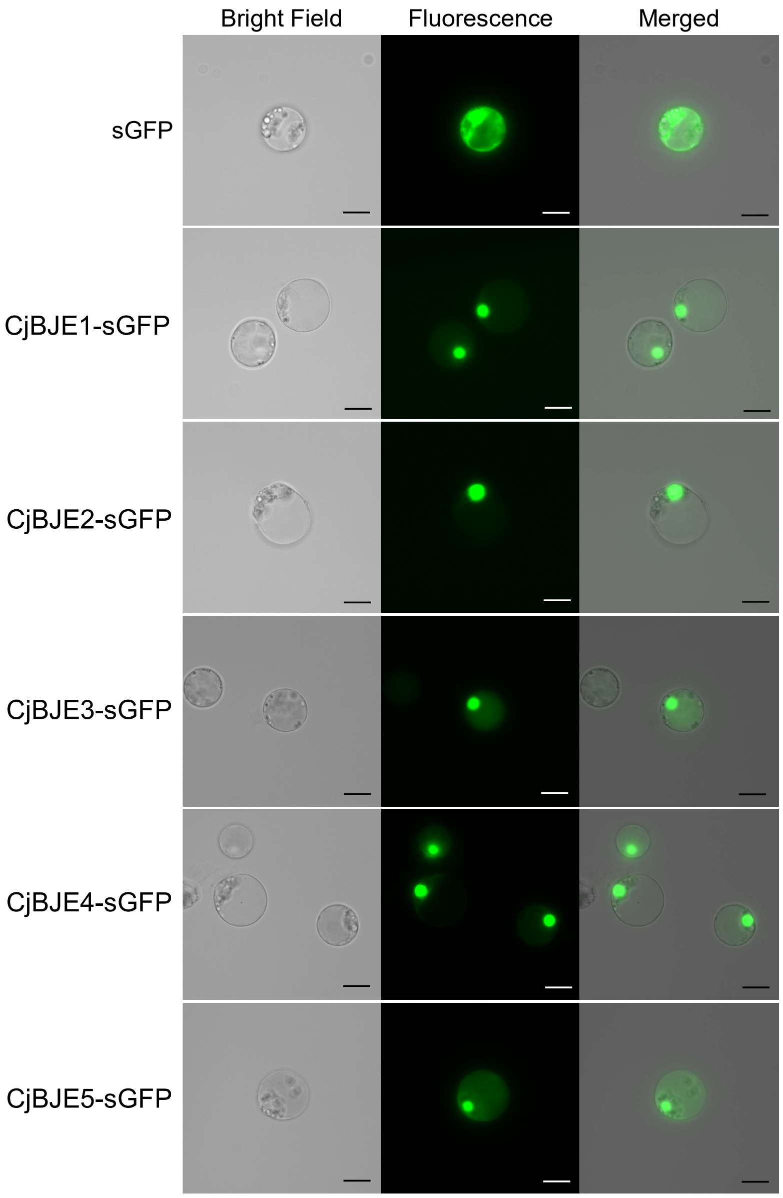


**Supplementary Figure S5. Subcellular localization of CjBJE proteins in *Coptis japonica* protoplasts.**
Expression vectors for sGFP-fused CjBJE proteins were introduced into *C. japonica* protoplasts and green fluorescence was observed after 24 h of incubation. Scale bars : 20 μm.

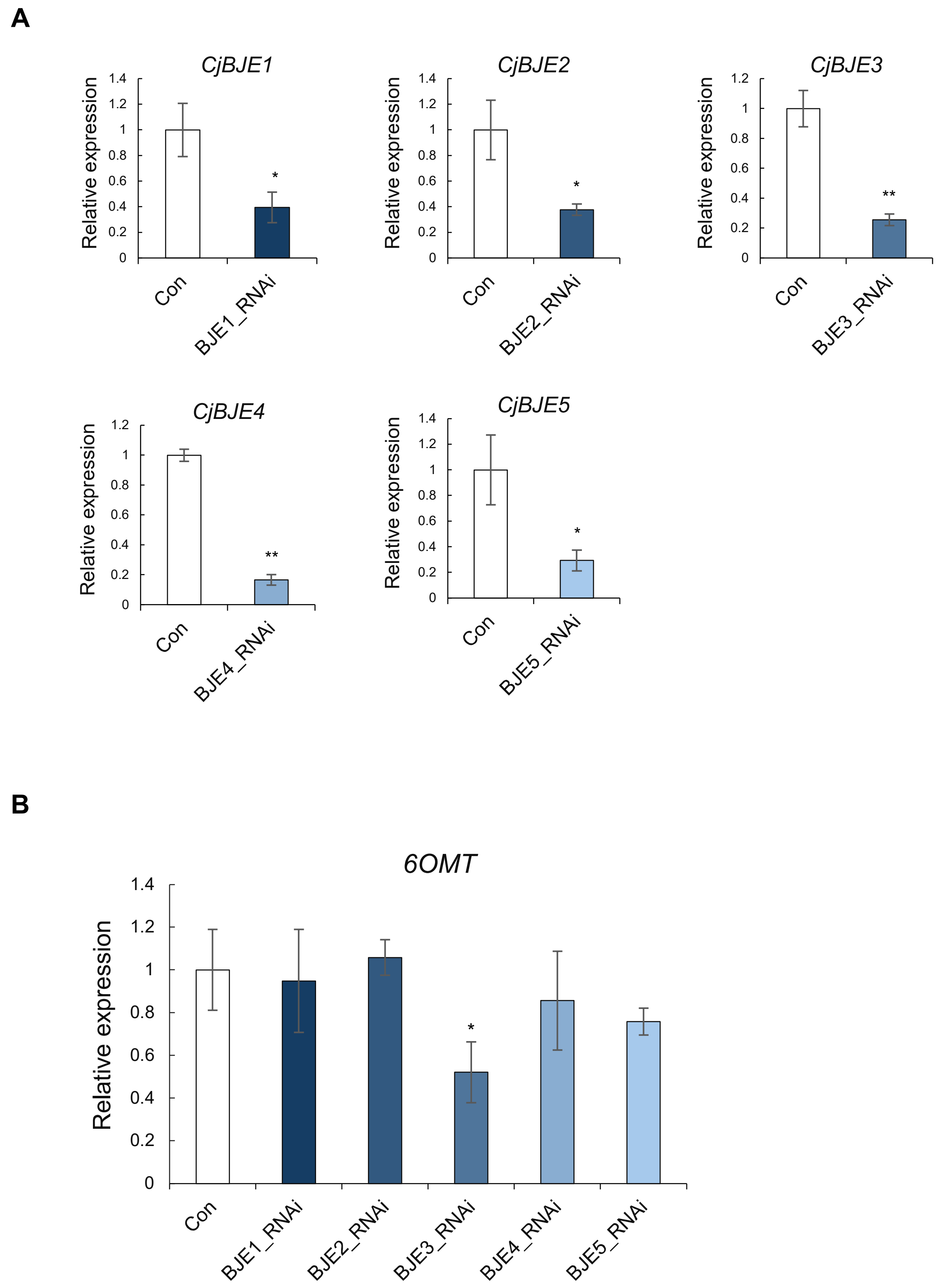


**Supplementary Figure S6. Transient RNA silencing of *CjBJE* genes in *Coptis japonica* protoplasts.**

Relative transcript levels of *CjBJE1–5* genes (A) and *Cj6OMT* (B) were determined by qRT-PCR and normalized to the *Cjα-tubulin* gene as an internal control. Relative expression levels are shown in terms of fold-change compared with control (Con) (set to 1.0). Error bars indicate standard deviation of three biological replicates. Asterisks indicate significant differences compared with control (Student's *t*-test: *$p < 0.05$, **$p < 0.01$).

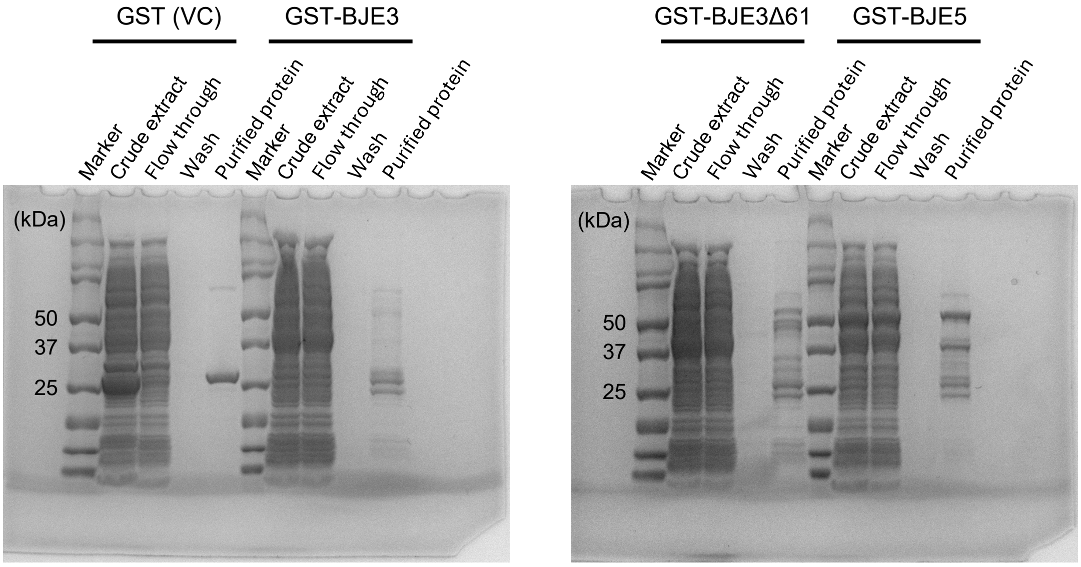


**Supplementary Figure S7. Coomassie brilliant blue-stained purified GST, GST-CjBJE3, GST-CjBJE3Δ61, and GST-CjBJE5 proteins.**

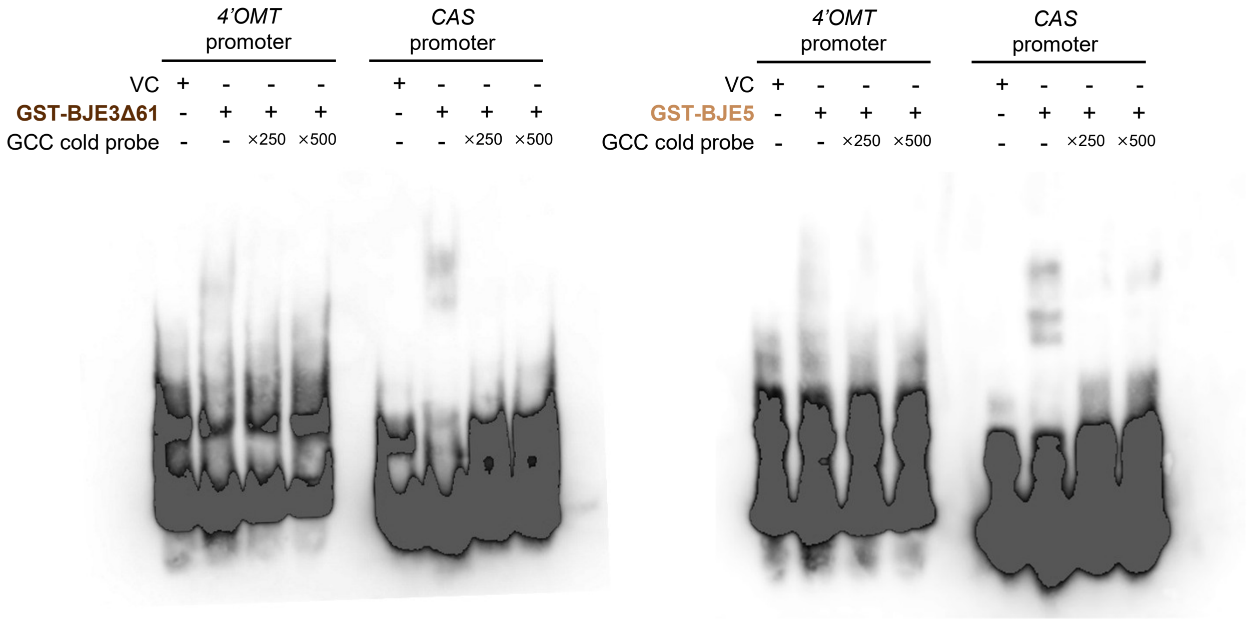


**Supplementary Figure S8. Electrophoresis mobility shift assay using GST-fused recombinant proteins**

Binding activity of truncated CjBJE3Δ61 and CjBJE5 recombinant proteins to biotin-labeled oligo DNA probes containing the GCC-like sequence. EMSA was performed using purified GST (VC) or CjBJE3Δ61 and CjBJE5. High amounts (250-fold and 500-fold) of non-labeled cold probes were used as competitors.

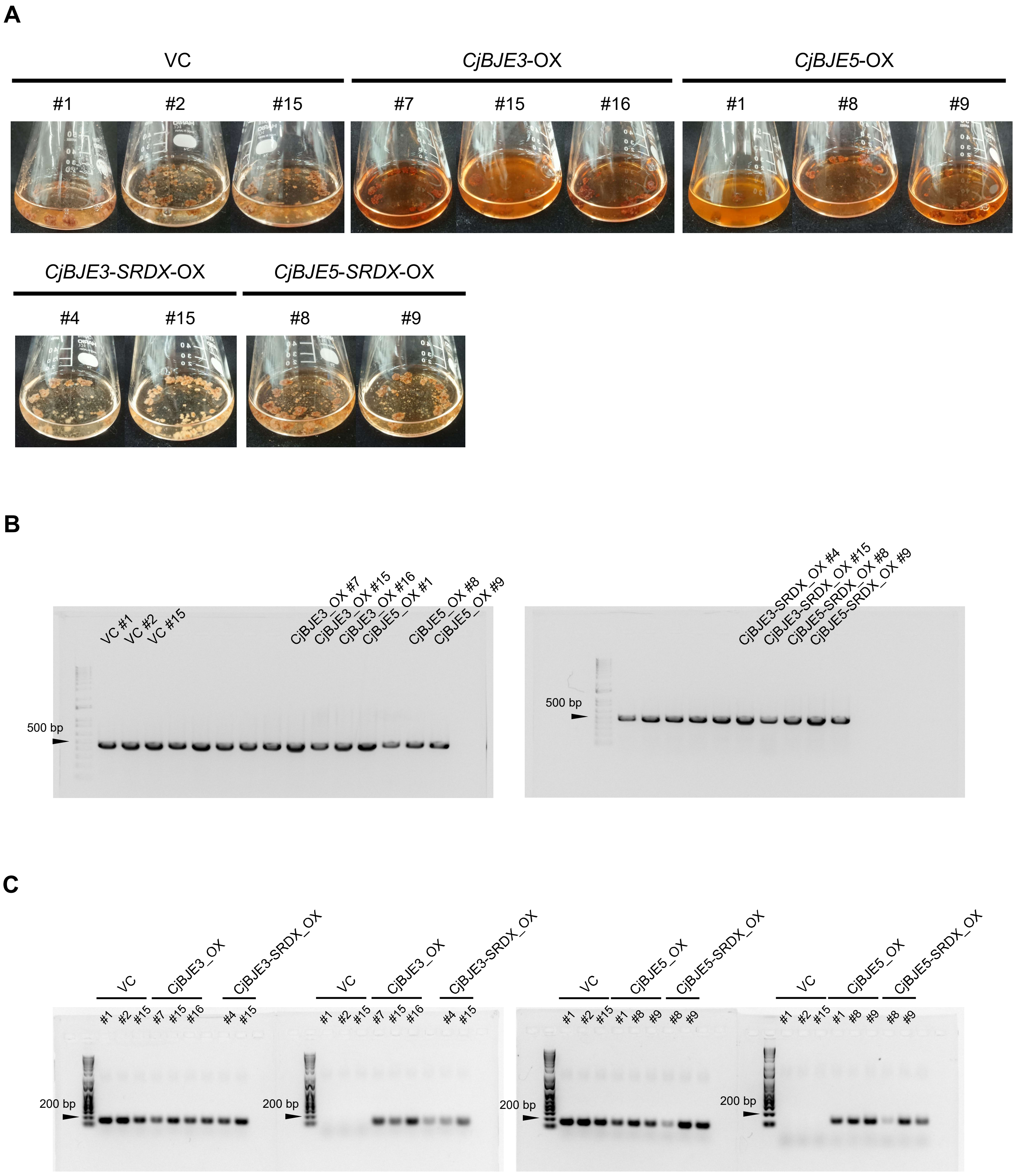


**Supplementary Figure S9. Establishment of transgenic cultured cell lines of the California poppy.**

(A) Three vector control (VC), *CjBJE3*-OX, and *CjBJE5*-OX lines, along with two *CjBJE3*_SRDX and *CjBJE5*_SRDX lines, were selected for transcriptome and metabolite analyses. (B) Detection of *NPTII* transgene in each cell line by genomic PCR. (C) Reverse transcription PCR showing heterologous expression of *CjBJE* genes in transgenic cultured California poppy cell lines using specific primer pairs constructed for qRT-PCR.

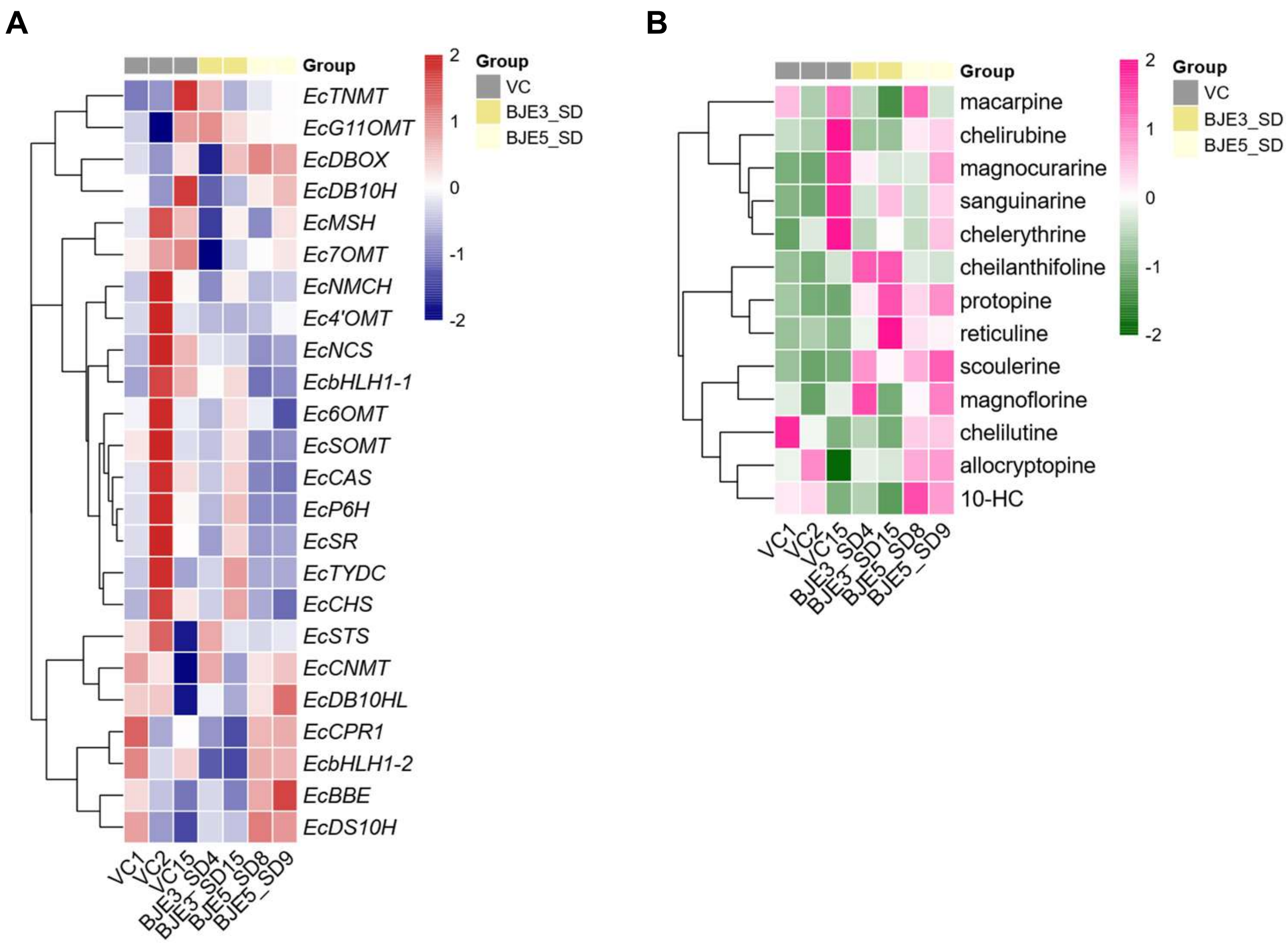


**Supplementary Figure S10. The overexpression of *CjBJE3*_SRDX and *CjBJE5*_SRDX in cultured *Eschscholzia californica* cells.**
(A) Gene expression profiling in both *CjBJE3*_SRDX and *CjBJE5*_SRDX cell lines. Heatmap and hierarchical clustering illustrate the expression patterns of known BIA biosynthetic genes. Transcript levels (TMM values) were normalized using Z-score transformation to visualize relative expression changes across samples. (B) Metabolite profiling in both *CjBJE3*_SRDX and *CjBJE5*_SRDX cell lines. Heatmap and hierarchical clustering show relative abundance of alkaloids. Data were standardized by calculating Z-scores for each metabolite across all samples.

**A**

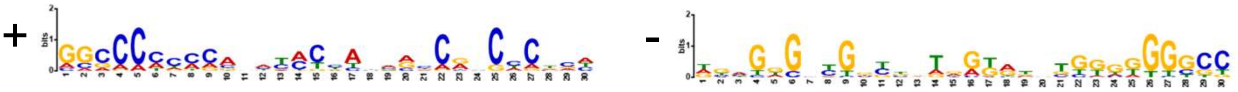

| | | | | | | |
|---|---|---|---|---|---|---|
| CYP76_L1 | + | 371 | 1.15e-13 | GTTTCGCTTG | GGGCCCCCGATATACTAGGGCCGGCCCTGA | TGATGGTGGA |
| OMT2pro | + | 1144 | 3.43e-12 | TTTTCGCTTT | AGGCCCCCCAACTACTTGGGCCGCCCCTGA | TTAGAATCAA |
| CYP82P5pro | - | 630 | 2.26e-10 | CTATATACCC | GCCCCCCCAAGCCCCCATGGAAGGGCCGAT | CAAATGGAAT |
| EcMSHpro | + | 428 | 3.25e-10 | GGGGCTCCCA | GCCCCACCGGGAGCCCAGCCCCGGGGCTCC | CGGGTTGGAG |
| EcBBE1pro | - | 1855 | 3.66e-10 | CTAATTCTAT | GGACCCCACATATCCTACAGTCATCTCACA | CGTGTAAGAT |
| EcSTSpro | - | 1332 | 4.12e-10 | ACTTCGCTTG | GGGCCTCCCAAATATTTGGGTCGGCCCTGT | GAATAAGCAC |
| NCS_L1pro | + | 174 | 9.28e-10 | GGCTTGGCCT | GGCCAACCCACTCACCCTAGCAGGCCATCA | TATGGCTGAC |
| CYP71_L3 | - | 1214 | 1.16e-9 | TGGGGGCCTG | CGCCCCCACGGGCCCCTACGTGGGCTCGCC | CCTGGCCTCA |
| BBE_L1pro | - | 1879 | 1.81e-9 | CAACTTCTAA | GGACCCCAGATATCCTATGAGCATCTCACA | CGTGTAGATA |
| EcBBE2pro | - | 1879 | 1.81e-9 | CAACTTCTAA | GGACCCCAGATATCCTATGAGCATCTCACA | CGTGTAGATA |
| CYP82P1v2pro | - | 338 | 2.26e-9 | TTCCGATATT | GGGCCCGCCCGACCTGATAAGCAACACTAA | GTACACGTAA |
| EcNMCHpro | - | 1731 | 5.27e-9 | ACGGGATTCC | GGGCACCCAAGGTCCACAAACCGTCTCTCT | GAAGTTTCCT |
| EcWRKY41pro | - | 94 | 1.93e-8 | TCAATTGGCT | GCACCAAACCGCCACCAAAACCGCCTCTTA | TAACTTTTAG |
| EcCPRpro | - | 1343 | 2.57e-8 | TCTTCCTCTC | ACCCACCACCTTTACCACAACCAGCACTTC | ATCTCATCAT |
| EcCNMTpro | - | 1540 | 6.44e-8 | GGTCGCACCA | GACCCCCCCTCAAACTACAATCTCCCCCGA | TCCCGTCCTC |
| EcSOMTpro | + | 392 | 7.69e-8 | TATTTCATTT | AGGCCTTCCAATTATTTGGACCGACCCTGC | ATATAAATAT |
| NMT1pro | - | 1415 | 9.17e-8 | AATTTACTTC | GGCACCCCAACCCATCATCCACAGCAAAAA | GTTTACAATG |
| CYP82N1pro | + | 899 | 1.19e-7 | CAGGTCGGTG | GACCCAGTCGGGTCGGATAGGCAACCCGCC | TATCCTAATA |
| EcWRKY36pro | + | 1 | 1.41e-7 | | GGCACCCCAATATTCAAACCCCACGCGTCC | TATCTTACCT |
| Ec4'OMTpro | + | 1799 | 1.41e-7 | CTTAACCTAA | ACCCCAAACCCACCCCTCAGCCATCCAAAT | GAAAATAAGA |
| CYP82N7pro | + | 1138 | 1.67e-7 | GACTTAGGGT | GGCGCATCCAAAGACCAAAGCCAGGTGGCT | CAGGACCAAC |
| EcDB10HLpro | + | 1220 | 2.14e-7 | CGGGAAACAT | GGCCCAACCAAAAATGCTAAACAACCCGAT | TTTTTGGGCT |
| EcDB10Hpro | - | 487 | 2.14e-7 | TTTCCTTAAT | GGCCCCCAACTTTGTGACTAGCTTCCCGCC | TGGCCCCCTA |
| BBE_L3pro | - | 116 | 2.33e-7 | TACGTGCATC | CGCCCTTGCAACCACCACCGCAGACACCAT | TTTAGGTGGC |
| CYP71_L2 | + | 1141 | 3.49e-7 | GTATATTAAA | GGCACCCAAGTCCCCTTGCATAGCCAATGA | TTGTATGGGC |
| DB10H_L2pro | - | 721 | 4.42e-7 | GGCCAAAAAC | GCCCCAGTCCGATAGAAACCACCCCCCCCC | CCCCCAGCCT |
| EcTYDCpro | + | 1297 | 5.58e-7 | TCCGACTACC | AGCACTTCCGATCATCATTACCACCACTCT | TTCTTTTCCT |
| CYP82N2v3pro | + | 682 | 7.56e-7 | ATCACTCCTC | CACCACACCACCTCCTCTCCACCTCACCCT | TCTTCTCCGG |

**B**

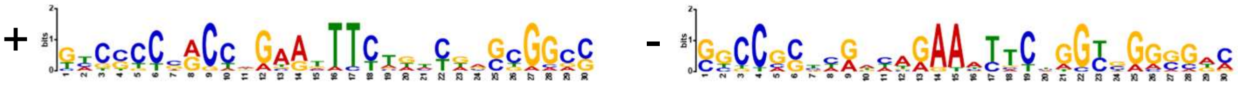

| | | | | | | |
|---|---|---|---|---|---|---|
| EcDB10Hpro | + | 313 | 2.61e-14 | AAATTTGAAG | GTCGCCGGCCGGAAATTTGGCCGGCCGGCG | ACCCATGTCG |
| BBE_L1pro | + | 1272 | 6.69e-14 | AGCATTGCAT | TACCCCCACCAGAAGTTCTGTTGAGCGGAC | TTGATGCATT |
| EcBBE2pro | + | 1272 | 6.69e-14 | AGCATTGCAT | TACCCCCACCAGAAGTTCTGTTGAGCGGAC | TTGATGCATT |
| EcCPRpro | - | 238 | 1.21e-12 | CTCCGATAAA | GTGGTCGACCAAAATTTCTGGCGACCGGCC | AGCAACCTTC |
| OMT3pro | + | 562 | 8.90e-12 | TTTTTTCATG | CTCGCCGGCCGGAAAATTTTCCGACCGGCG | ACCTAATGGT |
| CYP82P5pro | + | 539 | 6.84e-11 | CATTTCTGTT | GCCCCTCACATGCGTTTCTTACACGCGGGC | TGAGAACTCG |
| DB10H_L2pro | + | 650 | 2.38e-10 | GATTTCCGTT | GCCCCTCACTTGTGTTTCTTACATGCGGGC | TGAGAACAAG |
| CYP82N2v3pro | - | 733 | 8.03e-10 | TTTTAATAAG | GTGGTCGGCCGGAATTTTTAATTCCGGCCG | GCATGATTCC |
| CYP76_L1 | + | 1612 | 9.27e-10 | TGTCCGGCCA | GTCTCCCGGTTGAATTTCAGTCTGGTGACG | TGATCAGTCC |
| EcMSHpro | + | 379 | 2.78e-9 | GGGGCTGTGA | GCCCCCAGCCCGGGCTCCCATTTGGGAGCC | CGGGAGCATG |
| Ec6OMTpro | + | 1104 | 6.70e-9 | GTCTTTTTAG | TATCGCTACCAAAATTTCAATCAAGTGGCC | CACCTCTTGG |

**Supplementary Figure S11. MEME motif analysis of promoter regions in BIA biosynthetic genes.**

Promoter sequences (approximately 2,000 bp upstream of the transcription start site) were extracted for the BIA biosynthetic genes, as shown in Figure 8. Discriminative MEME motif discovery was performed by comparing upregulated genes in CjBJE3-OX and CjBJE5-OX cell lines with non-upregulated genes. Two representative GC-rich motifs, (A) and (B), were predicted from the top 10 enriched motifs.

>*CcBJE2* promoter

ACAATAATGCAGTAGTATAGACTTGATATCTAATCGAGGCTTTACACATGTTCATGTCCCTTGGCTTTCCTTCAAAATGGATACTATTTCTCAATT
TCCACAAAGATACCAAAGTACTAGCAACTGCCGAGTGCCAAAGATGTTTCAATCTTGGACTCCAAAGTCTTGCGACTCTAATAGCTTCAGTAAA
GCTATTGAAGTTCATAGAAATCTGAAATCTGCTTGTAGTCCAATGCCATAAAGCTCTGGAGAATGAACATTCCCAGAGTATATGTTGAAGGTTT
TCCACTTCAACCTTACATAATCTGCATGAAGAAGCAAGATGAACACCCTTTTTTTGAACCATACTATCAGTTGCCGCACAATTCTGAATGAGCTT
CCAAGCATTTGCCGCTAGTCTCGGGTGAATATGGGAGTTCCAGATACTTTTGCCCAACCAATTGGGTTGGCTTTCCTTCTTATGGAGTCGAAG
CGCAGTTTACAAGTGAATCTACCGTGAGTGTCCCGAGACCGCACCATATAATCTTCATGTTCGAGGCTCCTCTATGAACATTATCAATACTGCA
CCGGTTCCAACATATTTTGTAAAATAGGAGGAATGGACCAATTGTTGTTAACTAGGATAGAGTTAACCTTTGCATTAAAGCCCTTCAAAACAAG
GGAGAGTGAGTTCAAGAGATGTTTGAAGAGAGGTCGAAATGA**GCCGCC**TCGATCCTCTCGATTTTAGATCGGCTTTTGAAGATGCTTTGTAGT
ATTGACATTACCCTAGAGAATCACCCATTGGCTCCCACTAGATTTTACTTCTTAGGAGGCTTAAATCTTCTAATATCCGGTTTCCAAATTGAAGA
TGCTTTGTAGTATTGACAAGGGGTATCTCCATCCCTAGAGAAGTATTTCAAATTTTTGGTTCATGAATGTTGCAAAATCATCCACTTCAGTTAAA
ATTCTCCATGCAAGTTTCATTAGCATAGAAGAGTTAATATCTCGTAATCTTCTAATATCCGGTTTCAAATATTACCGTCATTGTTCAAAAGATAAT
TGTATAATAATATATGTGAGGGAGGGGTAAAACCATCTATTTGTATATTCAAATGCGTCTCCCTTGACATTGCCCTCAATTTTATAAACAATTAT
GAGATTATTGCCACTGATTGAAAGTTAAATTTTTGTGTTGTTAATCACACCACACGCATGGCAGACACCGCCATTTTAGGTATTTGTTACTGTCG
GACCCAATTTGTGTTAGGCGGATTCCAAAAAAAGCGATTATTACAAAAGAAAAAACAATTATTACAAAATATGCCATTTAAAAAAATTGAATTGAA
ACCCCAAACTTAAATTACCCGCCAGGTCGTTTTGGATTAAGTCTCAAATGATTTCATTGGTCCAACTTAGTCCAAACAATACGGCCATACCCCA
AAGGTCAAGAGCCAAAAAACGTAATTTGAATTTTTTTTTTCAAAGTTCAAATTCCAAACGCATACCCAAACCGTCGGATCGATTTAAAGTGTTAT
TATAGACTTAGCAAAACATTTTTAAAGCGTTCCAACTACCTGTACAGGCTGTACTCTGGGTTCATTAGCAAATTATATTCTTAACCGAAGTCATT
TAAAGTGTTATCCAAAGCGTTCCAACTGTTTTGTTTTAGCAAATTATTTCTTAACCGAAATTATCCGACAAACCGACAGGCATAGAAAAATTATT
GAGTGCTACGAGTGCTTTGTTTTTATTTTTATGCTATTCTTTCATTTATTTACTCGTGCATTTTATATTGTGCTTATCACAACGTACTCGAAAACA
GTTTAAAAAGGTATTGCAAAGCACAAGCAACAAAACAAAGTAGGATGTTTCATGTTTGCATCGTGGTAACTCCAAATCCATGACCAAACATCCC
AAGTTTCCAAACCCCTTTATATACCCTCCCTTCTATCACTCTATTTCTTAACTCTCCTCCTTCAGCTCTTTATTCTTTCTCTTTTGAATTCTGAGAC
TTCTACTCCA

>*CcBJE4* promoter

GTTGATATTGAAATCCACTTGACCAACTTTTTTTTTGCTAAAGCACTTGACCAACTAGATTTGCTCTGAAACAATAGAAATGGTTTCCAATACTATA
ATACATGAAATAGACACCTACATACGTTAAAATTTAGCTTCTATTCCAATTTTCGCCTTTTTTTTTGGGTATGTAAACTAAGAATACAACTTAGTT
GGCCTAGGACTATCTCTGGTAATGTGACCAGCTTGAACATTGTTTATTGATGCAATCACAAAATTACTTTTATTAAATATCAATAGAAAAATTATA
CAAAGATAGTACAGGACAATTGCACGATAAGTTGCACAGCTAGTCTTTTCTAAATAGCAAATTCCATTTTAGTGTCCATCATATTTTCCTAAATA
CATAGTTAATAGAGTCAGCTGGCCCTTATATTGTTGACAATTACCTTTTGCACCCTCTTCAAGAGTTAAGTTACTTGTGGTGCGGAAAATCTAAA
TGTACCAAATGCCCCTTCAAGTTTTTTTTTTTTTTTTTTTTTTTTTTTTTTTTTTTTTTTTTTTTTTTTTAACGAATTTTACAAATAATTATGATAAATATACAACAA
ACTGCGTCGCAAGTCCTTTTTAAATAGCAAATTATATCCTCCTAAATACAAACTCGTTAGAACAAATTGTCATTACGTTACTATACATAAACTTTT
TATTTATTTATTTTTTAAGTATGCAAAACTCTTGTAAAGTAAAGTGAAAATGCTGGTGTGCACAATTGATTCTTTACATATTTGACAAAAAATTATT
GCCTTTCGTTTGAAACCCGTTTAATTTTCAATCAACAACAATGGATGTCATGCGCGCCGATCATTTTGATGAGCATTTTTCAGTCAATAATTTGA
ATCATGAGATGCAATTTAAATTTTATTATGATCTCAAACTAACCATAACTAATATTATGTAATGTTTTTTTTTTGGTAACCATGTAATGTAGTATGT
AATGCAAGGCCTAAAATATTACCGATATATTCGATATATCACCGATATATATATCGTTTTTTTGGTATACCGATATCAATCTATCCGATATATATC
CACCGAATATTGAATCTTTTTTTTCCGATATATATCAGAATTTATGATATAAAACTTTATATCGCCGATATAATCGATATATGTGATATTTATGATA
TATATCGTTTGATTTTTTACCATTTTTTAATGTAAATATCTGATATTTTACAATATTTTATGATATTTTTTTACCGATATTTTTTATATTTTAGATAGTT
TGGAATAGTCACAAAAAAATAGATATATCTGATATTTCTAATATATGTTGGTTTTGGGTACACCAAAATATTCACCGAATTGGATATGTTGGCCTT
GATGTAATGTACTTAAAAAAAGAAATCGATATGTTGGGCTTGATGTAATGTACTTACAAAAAGAAATGTAGAATGTGCGTTCATGCATTTTACCC
ATAAACATCAGTGTCAAAACCTACAAGCAATTGGAGCCGGGCCAAACTAAACCAGGGGGAGGTCAGCCTTCTAGATACAATTATCCACAACAG
TTTAACACCCTTCCTGGCCATTCCAAATCCCATTCAACTGCCAAATCAATCCATATGCCACATTAGCAACGTACAAAAGAGGACGTCACCAGTA
CCTTGACCAG**GCCGCC**ATGAGTCAAATTGACCCACGTGCAAATAAATTACCCAAATATCAATAATATGTACTAGGCGTAGGACCACAGAGTAC
TAAAAACAGTGCTGAAACATCCTCCTTTGCCAATATTCCACTCTCATAGTCTCATTTCTTTATGCTTCACTTTGGTATTTTTTACCCTTAAAATCCT
CCCTCTCATTTGCTTATATAAAGGCTATTGCTTCAAATTTGTCAAAGCAACATAGACACCTAAGGAAAAAAATTTGCTAAACAAAAGTATCCCAA
TTTTAATACAATTCCATTCAAGGCAAACAACAATTCCAATTCATAGATGGAGACTTATTTTCTTCGATCAAGAAGTTTCGAT

>*CcBJE5* promoter

GCTTTTGAGTCATTAGCTATCTATATAGCAAGTACGGCCATAGGCCGGGCCAAATGGCTAGTATATTATAATAATGTGGTAGCAAAACGGGTTT
AAATTTCTATCTTGTTGAGGTTCAGTGATCCATTGTAAGGACCACTATGGTTGATTAATTACTACTTAATCCTCGTTCTATTGTTCAAGTTGTTTA
AGGTTGATTGATTAAGTAGTAATGATCTTATGTTGTAGTTTGTCACGTTTAGATACCATCAAAGAAATTTGTAATGATCATGAATCTTATTTTGTA
TTTTGCAGGTTCATGTACCAAATTTGGTTGGGTGGTGAAAATGATGTGTTTTTATAGTGAACTAATCAAGAATAATTTGGTTGTATTTTTGTTTCT
TATCCTAAATGATCCAATAGAAGGTTTCAGACCAATATCCAGATCTTATGAAACAACTTATAAGTGGTTATCTAAAAAAAAAAATGATGAACTTATA
AATTACTGAAGACATGCTGAATAAGTGCAAGTGCGCCCAGTCACTAGTGCAGCTTTCTATAGAAAGCGGCAGGGGCTAACGTTAAATACTTAA
ATGGTATATATCTGGATTAAGATACCATCTGAATTTATTATGTAGTGAAGGGGCCACTGATTGAATCAAGTTACAAGCAAGACCAACTCCTTTGA
ATCTTGAACATTAGACCATTATCAACGCACTGCTATATTTTTTTTTAAGACTTCAACGTTTTTAGTCTATCAAAATTTCTAAAAACGAGTCAAGTAC
CAATATATTTGAACAAAATTTGGAGTTTTTATTTTGGATGGGAATAGTTGTTCCATTCCAAACTGTAGAAAAGTCCAACGGGTACAAAATCGAGC
TGTTATAATTTTTTATTAAAAAAAAAGGCGGAAAAATCTTGGCCCCACAAAACTTGTCCCATTATAATTTTTTATTAAAAAAAAAGAAAAAAATACT
GGACGGCAAAACTTGTCCCGTCCCACTCTTGGACGGGACAACTTGTCCCGTGCAACTCATTTATGTTTTTCTCAGTATTTTAGTTTGAAATTTG
ACAATTTTTATAATCCAAATTGTGTTTTGGCTACGCGTTGGGATCAAAATTGCCAAATTCATTTCCATCTGTCACATTAGAAATGGGATTTGACA
ATGCGTTGATGATGCTCTTACTAACGTTATAACAAACAACCCCATTTAAATACTGAGCACCTGTGCTCTGAGTAATTGAATATATATATATAGAA
TAAAGAGTGAGCTCCTGTGCTATGAACAGTATGAAGCATATATGAGTAAAGAATAAAGGGGTACTGTAGATGTTCATTCCAATTGTTTTGAGGG
AAAAAAAATTCTTCCAGTTCAACAACTACTCCGAGATAGAGATATACTAATGGAAAAAGAAGGTTCTTTCCAAAAATAGTAAATAATAAAAGTTG
GAATAGTATAGTGCTATGAACAAAACCAATTTAAAAAAGCAATATAAGTTCTATTTTTCCTGATATGGGTACATAACTTGATAACTATATTGATAA
GTTACCATGGGTTATTGCCCGTTCAAGCATTACCCTTGATTACAAAATAGGGAGTTTTAACGTTTTGTCCGTGATGTTGAGGCAAAACCTCAAC
TTGGCACAGGTAGTTGCTATGGGGGCAGACTCAAACTAACAAATTGATTAAACAAAAGGAGGTCAGCATGCAGCATGCTACTTCCAAAAACAA
GCAATTTATTATG**GCCGCC**AGGGGTCGCAATTTACCCTCGTGCGACAATTTAGAGAGTATATGGTCCATTCCAGACCTCACTCAACATTATAGA
AACATCCTCCCTGGCCATTGTTCTTGCTTTGGTGGCCCTAGTCACACTCATTATTCTGACTTCTTTATCTCCTCTATAATTTTTCAAACTCTTCCC
TCTTCTTTGAGTATATATATTGGTCATACAGAATTTTAATAACAAAGTTTTTGTAAGCGTATAAGTGAAGAACCTTGGGAAACACTGCTAGAGTTT
AAAG

**Supplementary Figure S12. Nucleotide sequences of *Coptis chinensis BJE2*, *BJE4*, and *BJE5* homologous genes.**

Canonical GCC-box elements (GCCGCC) are highlighted in red.

**A**

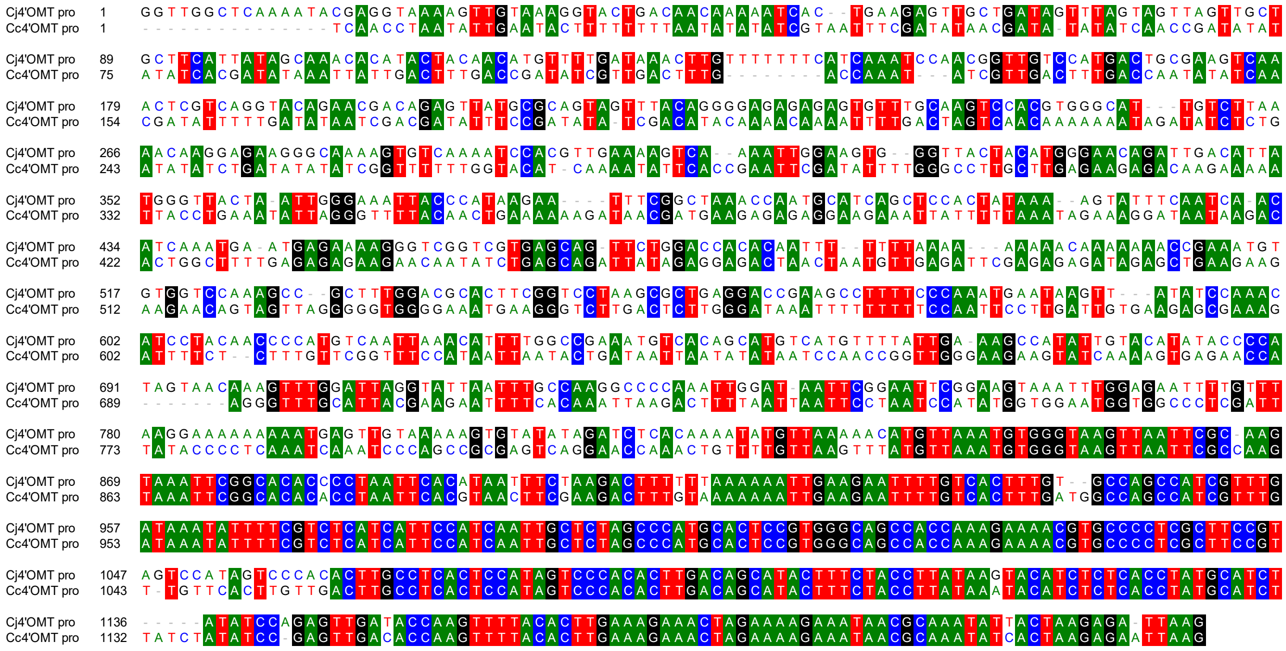


**B**

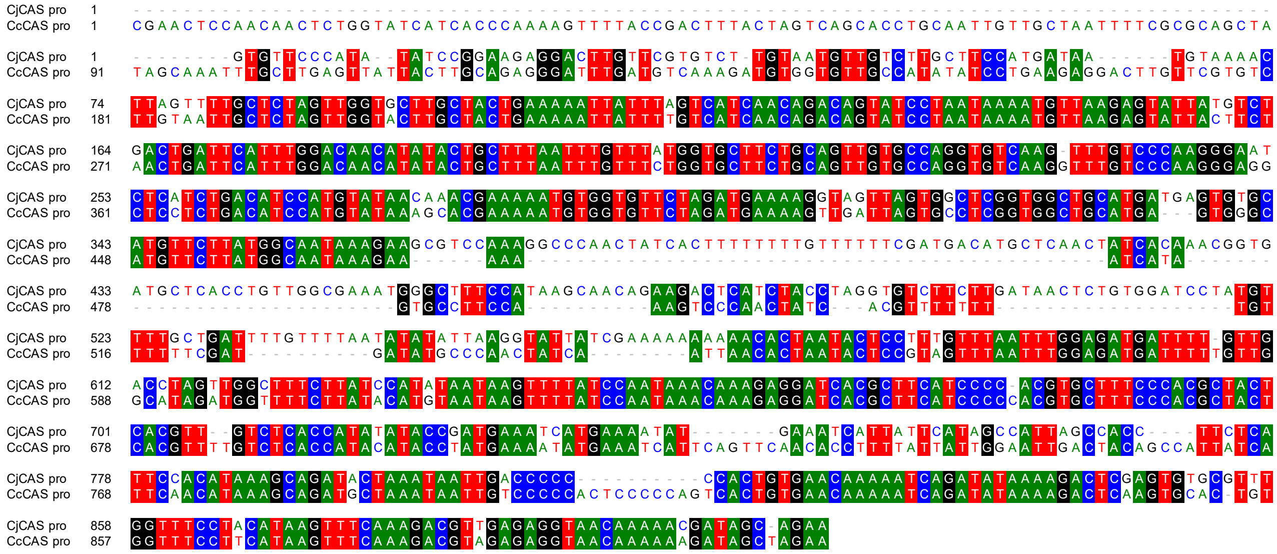


**Supplementary Figure S13. Nucleotide sequence alignment of *Coptis japonica* and *Coptis chinensis 4'OMT* and *CAS* promoter regions.**

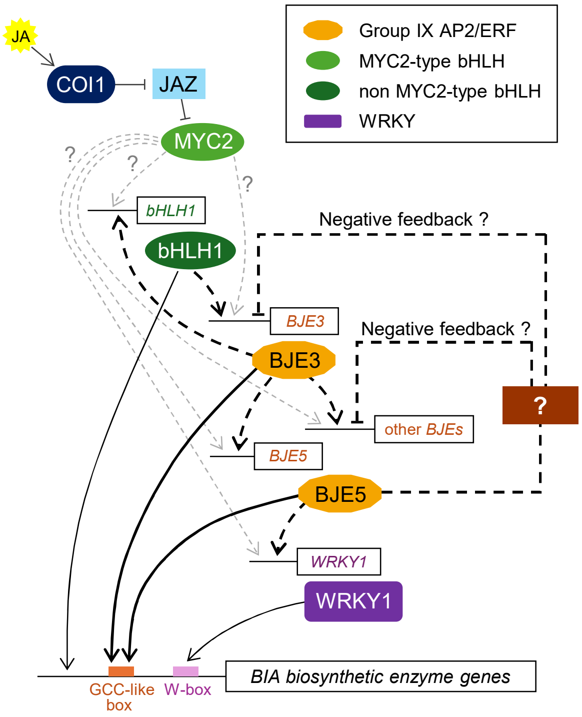


**Supplementary Figure S14. Proposed model for the transcriptional regulation of BIA biosynthesis mediated by CjbHLH1 and CjBJE transcription factors in *Coptis japonica*.** Expression of *CjbHLH1*, *CjWRKY1*, and *CjBJE* is induced by jasmonate, although whether CjMYC2 directly regulates these genes remains unclear. CjbHLH1 directly regulates several BIA biosynthetic enzyme genes by binding to their promoters. In the proposed regulatory network, CjbHLH1 regulates *CjBJE3* expression directly or indirectly. Subsequently, CjBJE3 regulates expression of *CjBJE5* and other *CjBJE* genes. Both CjBJE3 and CjBJE5 controlled expression of BIA biosynthetic enzyme genes by directly binding to GCC-like motifs in their promoter regions. Furthermore, CjBJE5 may indirectly regulate these biosynthetic genes by upregulating *CjWRKY1*. Our findings are also indicative of presence of a potential negative feedback loop involving CjBJE5, although precise mechanism remains unclear. Solid arrows indicate direct regulation, whereas dashed arrows represent regulatory steps that may be direct or indirect.

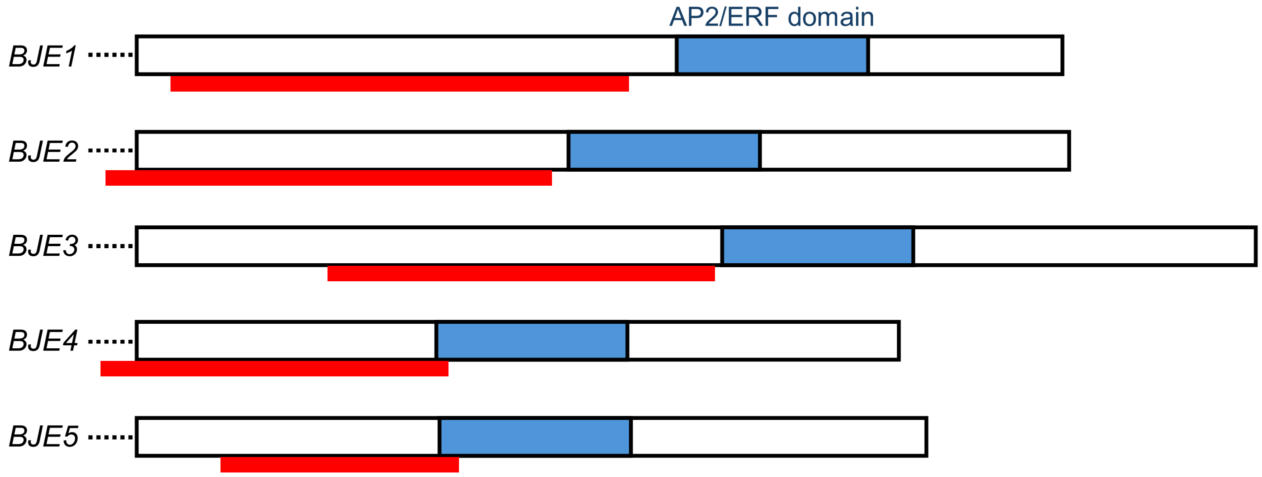


**Supplementary Figure S15. Target regions for transient RNAi in each *CjBJE* gene.** For dsRNA synthesis, partial regions indicated by red lines were amplified using primer pairs fused to T7 promoter sequence. White boxes represent coding sequences and AP2/ERF domains are indicated by blue boxes.

**Supplementary Table S1. Degenerate primer sequences for isolation of *AP2/ERF* genes.**

| Primer name | Oligonucleotide sequences (5' to 3') |
|---|---|
| ERFIX-consensus_Fw | TGGGGNAARTTYGCNGCNGARAT |
| ERFIX-degenerate_Fw1 | AARGGNAARCAYTAYMGNGGNGT |
| ERFIX-degenerate_Rv1 | CCNACNCKNARNGGRAARTT |
| ERFIX-degenerate_Fw2 | ACNATHGAYYWNATHMGNGARCA |
| ERFIX-degenerate_Rv2 | ATNGCYTTNSWNCCNCKCAT |
| ERFIX-degenerate_Fw3 | CCNTTYAAYRWNAAYGAYWSNGA |
| ERFIX-degenerate_Rv3 | ACNGGNSWRCANCCNTCYTC |

### Supplementary Table S2. Amino acid sequences of AP2/ERF domains.

| Name | Accession or Gene ID | Amino acid sequences |
|---|---|---|
| CjBJE1 | LC941495 | HYRGVRQRPWGKFAAEIRDPAKNGARVWLGTFETAEDAALAYDRAAYRMRGSRALLNFP |
| CjBJE2 | LC941496 | HYRGVRRRPWGKYAAEIRDSNRQGSRLWLGTFETAIEAAKAYDRAAFNMRGSKAILNFP |
| CjBJE3 | LC941497 | HYRGVRQRPWGKFAAEIRDPSRKGARVWLGTFETAIEAARAYDRAAFKMRGSKAILNFP |
| CjBJE4 | LC941498 | YRGVRRRPWGKFAAEIRDSTRHGIRVWLGTFDSAEAAALAYDQAAFSMRGSMAVLNFP |
| CjBJE5 | LC941499 | YRGVRRRPWGKFAAEIRDSTRHGVRVWLGTFDSAESAALAYDQAAFSMRGPMAVLNFP |
| AT4G17500 AtERF1 | O80337.2 | HYRGVRQRPWGKFAAEIRDPAKNGARVWLGTFETAEDAALAYDRAAFRMRGSRALLNFP |
| AT5G47230 AtERF5 | O80341.1 | HYRGVRQRPWGKFAAEIRDPNKRGSRVWLGTFDTAIEAARAYDEAAFRLRGSKAILNFP |
| AT3G23240 ERF1 | Q8LDC8.2 | SYRGVRRRPWGKFAAEIRDSTRNGIRVWLGTFESAEEAALAYDQAAFSMRGSSAILNFS |
| ORCA2 | CAB93939.1 | RYRGVRRRPWGKFAAEIRDPKRKGSRIWLGTYETAEDAALAFDQAAFQLRGSRARLNFP |
| ORCA3 | ABW77571.1 | RYKGVRRRPWGKFAAEIRDPKKKGSRIWLGTYETPEDAALAYDAAAFNMRGAKARLNFP |
| ORCA4 | QTJ02267.1 | RYRGVRRRLWGKFAAEIRDPNKKGSRIWLGTYETPEDAALAYDGAAFKMRGSKAKLNFP |
| ORCA5 | QTJ02266.1 | RYRGVRRRPWGRFAAEIRDPNKKGSRIWLGTYETPEDAALAYDRAAFQIRGAKALLNFP |
| ORCA6 | QTJ02268.1 | RYRGVRRRPWGTFAAEIRNPMKKGSRIWLGTYETPEDAAIAYDRAAFKLRGAKARLNFP |
| GAME9/JRE4 | XP_015164502.1 | RFIGVRRRQWGTFSAEIRDPNRRGARLWLGTYESPRDAALAYDQAAYKIRGTKVRLNFP |
| NtERF189 | BAN57618.1 | RYIGVKRRPWGTFSAETRDPSRKGEGARLWLGTYETAEDAALAYDQAAFKIRGSRARLNFP |
| NtORC1 | NP_001312507.1 | RYIGVKRRPWGTFSAEIRDPERRGARLWLGTYETPEDAALAYDQAAFKIRGSRARLNFP |
| NtERF32 | NP_001311965.1 | HYRGVRQRPWGKFAAEIRDPAKNGARVWLGTYETAEEAALAYDKAAYRMRGSKALLNFP |
| OpERF2 | LC171328 | RYRGVRRRPWGKFAAEIRDPKKRGSRIWLGTYETPEDAALAYDGAAFRMRGARAMLNFP |
| AaERF1 | AEQ93554.1 | HYRGVRQRPWGKFAAEIRDPAKNGARVWLGTYETAEEAATAYDIAAYRMRGSKALLNFP |
| AaERF2 | AEQ93555.1 | HYRGVRQRPWGKFAAEIRDPNKKGTRVWLGTFETAIEAAKAYDRAAFKLRGSKAILNFP |
| CrERF5 | MK862158 | HYRGVRQRPWGKFAAEIRDPNKKGSRVWLGTFDTAVEAAKAYDRAAFRLRGSKAILNFP |
| AaORA | AGB07586.1 | RYRGVRRRPWGKFTAEIRNPEKKKARLWLGTFDTPEQAALAYDRAAFKFHGSRAKVNFP |
| EcAP2/ERF1 | Eca_sc000448.1_g0520.1 | HYRGVRQRPWGKFAAEIRDPAKNGARVWLGTFETAEDAALAYDRAAYRMRGSRALLNFP |
| EcAP2/ERF2 | Eca_sc006292.1_g0200.1 | KYRGVRRRPWGKFAAEIRDPSRKGSRVWLGTFETAIDAAKAYDSAAFKMRGSKAILNFP |
| EcAP2/ERF3 | Eca_sc006292.1_g0150.1 | KYRGVRQRPWGKFAAEIRDPNRKGTRVWLGTYETAIDAARAYDRAAFKMRGSKAIVNFP |
| EcAP2/ERF4 | Eca_sc006292.1_g0190.1 | KYRGVRQRPWGKFAAEIRDPSRGGSRVWLGTFDTAIDAAKAYDRAAFKMRGSKAICNFP |
| EcAP2/ERF6 | Eca_sc000360.1_g1660.1 | HYRGVRRRPWGKFAAEIRDSARQGARVWLGTFDTAEEAALAYDKAAFRMRGSKALLNFP |
| EcAP2/ERF12 | Eca_sc194641.1_g1370.1 | AYRGVRTRPWGKFAAEIRDSTRNGVRVWIGTFDTAEEAALAYDQAAFSMRGSMAILNFS |
| EcAP2/ERF16 | Eca_sc183659.1_g1650.1 | HFRGVRRRPWGKYAAEIRDSSRKGARVWLGTFDTAEKAALAYDKAALKMRGPRTYLNFP |
| EcAP2/ERF17 | Eca_sc194316.1_g0160.1 | AYRGVRKRPWGKFAAEIRDSTRHGIRVWLGTFDSAEAAALAYDQAAFSMRGSMAILNFP |
| EcAP2/ERF18 | Eca_sc004486.1_g0660.1 | HYRGVRERPWGKFAAEIRDSARQGARVWLGTFNTAEEAAMAYDRAAYKMRGAKALLNFT |
| AT4G18450 AtERF091 | NP_193580.1 | KYRGVRKRPWGKFAAEIRDSTRNGVRVWLGTFQTAEEAAMAYDKAAVRIRGTQKAHTNFQ |
| AT2G31230 AtERF15 | AEC08512.1 | SYRGVRKRPWGKFAAEIRDSTRNGIRVWLGTFDKAEEAALAYDQAAFATKGSLATLNFP |
| AT5G43410 AtERF096 | AED94958.1 | KYRGVRRRPWGKYAAEIRDSRKHGERVWLGTFDTAEEAARAYDQAAYSMRGQAAILNFP |
| AT3G23230 AtTDR1 | AEE76737.1 | RFRGVRRRPWGKFAAEIRDPSRNGARLWLGTFETAEEAARAYDRAAFNLRGHLAILNFP |
| AT2G44840 AtERF13 | AEC10473.1 | QYRGVRRRPWGKFAAEIRDPKKNGARVWLGTYETPEDAAVAYDRAAFQLRGSKAKLNFP |
| AT5G61600 AtERF104 | AED97495.1 | HYRGVRRRPWGKYAAEIRDPNKKGCRIWLGTYDTAVEAGRAYDQAAFQLRGRKAILNFP |
| AT5G07580 AtERF106 | AED91182.2 | HYRGVRRRPWGKFAAEIRDPAKKGSRIWLGTFESDVDAARAYDCAAFKLRGRKAVLNFP |
| AT1G06160 ORA59 | Q9LND1.1 | SYRGVRKRPWGKFAAEIRDSTRKGIRVWLGTFDTAEAAALAYDQAAFALKGSLAVLNFP |

**Supplementary Table S3. Primer sequences for qRT-PCR.**

| Primer name | Oligonucleotide sequences (5' to 3') |
|---|---|
| CjBJE1_RT_Fw | ACGTTCATCACCTGAGCCTACG |
| CjBJE1_RT_Rv | CCAACTCACTGCCTGAACCACT |
| CjBJE2_RT_Fw | TGCAGAAGAAGAGAAGGCAGTGA |
| CjBJE2_RT_Rv | AGGCTCTCCCAAATGTCAGTCC |
| CjBJE3_RT_Fw | CCATTAACACCGTCGAGTTGGA |
| CjBJE3_RT_Rv | AGATAACGGGGACAGTGGAGGA |
| CjBJE4_RT_Fw | TGTGGGTTTGAGGATGGATGTT |
| CjBJE4_RT_Rv | GCTGGCTTTCTTGTTCGTGGAT |
| CjBJE5_RT_Fw | CGCGAGTAGAGAATGTGGTGGA |
| CjBJE5_RT_Rv | AGCAAGAACTGGAGCTGGGACT |
| CjTYDC_RT_Fw | AATGAAACACGTGCCAATGA |
| CjTYDC_RT_Rv | AAACAACGTGTCGCTCCTCT |
| CjNCS (DOX)_RT_Fw | GACTGGGGCTTCTTCCAGTT |
| CjNCS (DOX)_RT_Rv | TGGCCATAACCTTCCATACC |
| CjNCS2 (PR10)_RT_Fw | CGATGAAGTGTGGAGTGTCG |
| CjNCS2 (PR10)_RT_Rv | CCTTCACCTCCATCACCAGT |
| Cj6OMT_RT_Fw | GTGCATCCTTCACGACTGG |
| Cj6OMT_RT_Rv | TGCATCATGGATGAGCTTCT |
| CjCNMT_RT_Fw | ACCACACATGAGATGGCTGA |
| CjCNMT_RT_Rv | CCCAGCAGGAAACACGTACT |
| CjNMCH_RT_Fw | GAGGTTTTTGAGTTCTGATGTGG |
| CjNMCH_RT_Rv | GGACAATGAGGAGAGGTGGA |
| Cj4'OMT_RT_Fw | GGAAGGACACCCTGATCAAA |
| Cj4'OMT_RT_Rv | TTCCTCCACCAACATCAACA |
| CjBBE_RT_Fw | GACGAAGCACACCAGTGAAA |
| CjBBE_RT_Rv | AACGTGAGCCAAAATCCATC |
| CjSMT_RT_Fw | GGATTTCTTTCAGTATGCTGG |
| CjSMT_RT_Rv | TCTCTATCCCGTCTCCCAATC |
| CjCAS_RT_Fw | TGGTGAGGCCACTTCTCTCT |
| CjCAS_RT_Rv | TCTTGTGCTCCTTGTTCACG |
| CjTHBO_RT_Fw | CCAATGCCTTGATCTTCGTT |
| CjTHBO_RT_Rv | CAAAGTCATGACCCCCACTT |
| CjAlpha(α)-tubulin_RT_Fw | CAGTGAAACTGGTGCTGGAAAG |
| CjAlpha(α)-tubulin_RT_Rv | ATGAGCTGTTCTGGGTGAAACA |
| CjActin_RT_Fw | GTCACACCGTCCCCATTTA |
| CjActin_RT_Rv | GTCACGGACGATTTCTCGTT |
| CjbHLH1_RT_Fw | TGCTTCCTCGGTTGCTATCT |
| CjbHLH1_RT_Rv | TGCATCTATTGGTGCTCCTG |
| CjWRKY1_RT_Fw | TGAGCATGCACTCCCTCATA |
| CjWRKY1_RT_Rv | TGGAGGAATATGGGCAAAA |
| CjATPase_RT_Fw | TCAACAGCCAAAGTTGTTCG |
| CjATPase_RT_Rv | AATTCAGTCTGCCCGTGATT |
| CjCM_RT_Fw | TTTAATCCGCCAAGAGGACA |
| CjCM_RT_Rv | CCATGAAACCCATCCATAGC |

**Supplementary Table S4. Primer sequences for synthesis of dsRNA.**

| Primer name | Oligonucleotide sequences (5' to 3') |
|---|---|
| CjBJE1-T7-Fw | TAATACGACTCACTATAGGGAGACCACATTCAGTTCGTCGACACTTACTCG |
| CjBJE1-T7-Rv | TAATACGACTCACTATAGGGAGACCACCTTTTCTGGTAGTCGAGCCAGATT |
| CjBJE2-T7-Fw | TAATACGACTCACTATAGGGAGACCACTGAAACTTCTACTCCAATGGCAAC |
| CjBJE2-T7-Rv | TAATACGACTCACTATAGGGAGACCACACGACTTCTGTTCTGAAACACTGG |
| CjBJE3-T7-Fw | TAATACGACTCACTATAGGGAGACCACCTGCACCAATTTACTCTCAAGCAT |
| CjBJE3-T7-Rv | TAATACGACTCACTATAGGGAGACCACCAGAATCTACTGGTTTCGATGCAG |
| CjBJE4-T7-Fw | TAATACGACTCACTATAGGGAGACCACGCAACATAGACGCCTAAGAACAAA |
| CjBJE4-T7-Rv | TAATACGACTCACTATAGGGAGACCACCGCACGCCTCTATATGCTTTATCT |
| CjBJE5-T7-Fw | TAATACGACTCACTATAGGGAGACCACTCCATTCGATGACTTGGCTACTAA |
| CjBJE5-T7-Rv | TAATACGACTCACTATAGGGAGACCACGCCTTCGGACACCTCTATAAGACT |

**Supplementary Table S5. Oligonucleotide sequences of EMSA probes.**

| Probe name | Oligonucleotide sequences (5' to 3') |
|---|---|
| 4'OMTpro-212-165_s | CTCCGTGGGCAGCCACCAAAGAAAACGTGCCCCTCGCTTCCGTAGTCC |
| 4'OMTpro-212-165_as | GGACTACGGAAGCGAGGGGCACGTTTTCTTTGGTGGCTGCCCACGGAG |
| CASpro-156-121_s | CATAGCCATTAGCCACCTTCTCATTCCACATAAAGCA |
| CASpro-156-121_as | TGCTTTATGTGGAATGAGAAGGTGGCTAATGGCTATG |

**Supplementary Table S6. UPLC-MS analytical parameters and identification criteria for targeted metabolites.**

| Compound | Formula | Adduct | SIR target *m/z* | RT (min) | Reference/Basis |
|---|---|---|---|---|---|
| reticuline | $C_{19}H_{23}NO_4$ | $[M+H]^+$ | 330.2 | 3.35 | authentic standard |
| scoulerine | $C_{19}H_{21}NO_4$ | $[M+H]^+$ | 328.2 | 3.72 | authentic standard |
| cheilanthifoline | $C_{19}H_{19}NO_4$ | $[M+H]^+$ | 326.1 | 4.12 | authentic standard |
| protopine | $C_{20}H_{19}NO_5$ | $[M+H]^+$ | 354.1 | 5.23 | authentic standard |
| allocryptopine | $C_{21}H_{23}NO_5$ | $[M+H]^+$ | 370.2 | 5.75 | authentic standard |
| magnocurarine | $C_{19}H_{24}NO_3^+$ | $[M]^+$ | 314.2 | 2.35 | authentic standard |
| magnoflorine | $C_{20}H_{24}NO_4^+$ | $[M]^+$ | 342.2 | 3.08 | authentic standard |
| sanguinarine | $C_{20}H_{14}NO_4^+$ | $[M]^+$ | 332.1 | 6.51 | authentic standard |
| chelerythrine | $C_{21}H_{18}NO_4^+$ | $[M]^+$ | 348.1 | 7.63 | authentic standard |
| 10-hydroxychelerythrine | $C_{21}H_{18}NO_5^+$ | $[M]^+$ | 364.1 | 7.45 | Takemura et al., 2010 |
| chelirubine | $C_{21}H_{16}NO_5^+$ | $[M]^+$ | 362.1 | 7.96 | Takemura et al., 2010 |
| chelilutine | $C_{22}H_{20}NO_5^+$ | $[M]^+$ | 378.1 | 8.89 | Takemura et al., 2010 |
| macarpine | $C_{22}H_{18}NO_6^+$ | $[M]^+$ | 392.1 | 9.39 | Takemura et al., 2010 |
